# Computational Analysis of Therapeutic Neuroadaptation to Chronic Antidepressant in a Model of the Monoaminergic Neurotransmitter and Stress Hormone Systems

**DOI:** 10.1101/633990

**Authors:** Mariam Bonyadi Camacho, Warut D. Vijitbenjaronk, Thomas J Anastasio

## Abstract

The clinical practice of selective serotonin reuptake inhibitor (SSRI) augmentation relies heavily on clinical judgment and trial-and-error. Unfortunately, the drug combinations prescribed today fail to provide relief for all treatment-resistant depressed patients. In order to identify potentially more effective treatments, we developed a computational model of the monoaminergic neurotransmitter and stress-steroid systems that neuroadapts to chronic administration of combinations of antidepressant drugs and hormones by adjusting the strengths of its transmitter-system components (TSCs). We used the model to screen 60 chronically administered drug/hormone pairs and triples, and identified as potentially therapeutic those combinations that raised the monoamines (serotonin, norepinephrine, and dopamine) but lowered cortisol following neuroadaptation in the model. We also evaluated the contributions of individual and pairs of TSCs to therapeutic neuroadaptation with chronic SSRI using sensitivity, correlation, and linear temporal-logic analyses. All three approaches found that therapeutic neuroadaptation to chronic SSRI is an overdetermined process that depends on multiple TSCs, providing a potential explanation for the clinical finding that no single antidepressant regimen alleviates depressive symptoms in all patients.

## 1. Introduction

Depression is a debilitating psychological disorder and a leading cause of physical disability that lacks effective pharmacological interventions (Cipriani et al., 2016; Friedrich, 2017). The current first-line treatment for depression is chronic administration of selective serotonin reuptake inhibitors (SSRIs). This treatment provides complete relief from depressive symptomatology in only one-third of patients (Turner et al., 2008). SSRI non-responders are usually treated either by switching them to another SSRI, or by prescribing a second drug (such as an antipsychotic, like Asenapine, or an atypical antidepressant, like Bupropion) to augment SSRI action on the monoamines (Barowsky and Schwartz, 2006). Although these strategies have been shown to increase the proportion of depressed patients who respond to pharmacotherapy, an effective antidepressant combination unfortunately is never found in the 10-30% of depressed patients who are classified as treatment-resistant (Ananth, 1998; Barowsky and Schwartz, 2006).

The current approach to antidepressant drug design is based on the monoamine hypothesis. Mood is largely determined by the 3 monoaminergic neurotransmitters: serotonin (5HT), secreted from the dorsal raphe (DR); norepinephrine (NE), secreted from the locus coeruleus (LC); and dopamine (DA), secreted from the ventral tegmental area (VTA). The monoamine hypothesis was developed in the 1960s, following on clinical observations that drugs that elevate brain levels of 5HT, NE, or DA had mood-elevating effects (Schildkraut, 1965). The low efficacy-rate of SSRIs and of current SSRI augmentation strategies has led the field to investigate pharmacological targets beyond the monoamines that could improve antidepressant response efficacy, especially the interactions between the monoaminergic neurotransmitter and other neurotransmitter and hormone systems.

The most commonly observed risk factors for depression are stressful life events. For that reason, our current model is focused on the interactions between the monoaminergic neurotransmitter systems and the neuroendocrine response to stress (Kendler et al., 1999; Kendler and Gardner, 2016). Stress leads to activation of the hypothalamic-pituitary-adrenal (HPA) axis, resulting in elevated plasma levels of the stress-steroid, cortisol (Kendler et al., 1999). Briefly, HPA axis activation begins with corticotropin releasing factor (CRF) secretion from the paraventricular nucleus (PVN) of the hypothalamus in response to stress. Activation of pituitary gland CRF receptors by CRF results in adrencorticotropic hormone (ACTH) release from the pituitary gland. Binding of ACTH to its receptors on the adrenal gland promotes cortisol release from the adrenal cortex (Morris et al., 2012; Pariante and Lightman, 2008). In the short-term, cortisol release in response to HPA axis activation can be beneficial for dealing with stress (Dhabhar and Mcewen, 1997; McEwen, 2004). However, chronic elevations of blood cortisol levels are related to depressive symptomology, and response to antidepressant drugs is associated with normalization of plasma cortisol levels (Johnson et al., 1992; Wong et al., 2000).

The interactions between the monoaminergic neurotransmitter system and the HPA axis are complex. For example, activation of monoaminergic receptors on the PVN, pituitary gland, and adrenal gland by monoaminergic neurotransmitters has been found to enhance HPA axis activity (Dinan, 1996; Ma and Morilak, 2005; Ziegler et al., 1999), while cortisol can alter expression of proteins involved in monoaminergic synaptic transmission, including serotonergic receptors, monoamine synthesis enzymes, and monoamine oxidase (MAO) (Hucklebridge et al., 1998; McAllister-Williams et al., 2007; Nexon et al., 2011).

Here we represent the major interactions between the monoaminergic neurotransmitter systems and the HPA axis in a computational model that we refer to as the <u>M</u>onoamine-<u>S</u>tress model (MS-model). It extends our previously published computational model of the monoaminergic neurotransmitter system but differs in its structure, training procedure, and analysis (Camacho and Anastasio, 2017). Specifically, the MS-model takes the form of a recurrent network in order to use a more efficient learning procedure to train its more extensive representations of neurobiological interactions, and to conform to a larger set of experimental observations.

Acute administration of substances (such as drugs or hormones) has been observed to alter neuronal activity levels, and chronic (days to weeks) substance exposure can lead to adaptive changes in neurons that move their activity levels back toward their original levels (Blier and De Montigny, 1987; Turrigiano, 2008, 1999). We simulated neuroadaptive changes by allowing a subset of transmitter system components (TSCs, mainly proteins such as neurotransmitter or neurohormone receptors or transporters) to adjust their strengths (corresponding to factors such as expression levels, sensitivities, and synaptic locations) incrementally up or down. TSC-strength configurations that restored the activities of DR, LC, VTA, and PVN back toward normative baselines were referred to as “adapted.” We found that many different TSC-strength configurations could produce adaptation to chronic administration of any drug or drug combination, and each adapted configuration had its own, unique pattern of neurotransmitter and hormone levels.

Specific drug and hormone combinations were evaluated for their effects on 5 properties of interest, which were the level of adaptation, the levels of the 3 monoamines, and of level of cortisol. Therapeutic states were adapted states that also had elevated monoamine levels and reduced cortisol levels. By examining the distributions of monoamine levels over all of the adapted TSC-strength configurations, we identified drug and hormone combinations that could potentially be therapeutic for a larger proportion of depressed patients. We also constructed a heatmap of the average monoamine levels that could be expected in a patient population following adaptation to chronic administration of specific drug and/or hormone combinations. This heatmap could be used to guide treatment plans toward desired elevations in specific monoamines.

In addition to identifying and analyzing potential drug and hormone combinations of interest, we also determined whether specific TSCs contribute more than others to the establishment of a therapeutic state following adaptation to chronic administration of SSRI. Specifically, we used sensitivity and correlation analyses as well as linear temporal-logic to determine if specific TSCs or pairs of TSCs mediate therapeutic neuroadaptations. Our analyses determined that no single TSC or pair of TSCs are determinative of the therapeutic state, and that neuroadaptation to chronic antidepressant is a highly overdetermined process to which multiple TSCs can contribute in many different ways.

This analysis provides a possible explanation for the clinical finding that no single antidepressant drug or combination alleviates depressive symptoms in all patients (Barowsky and Schwartz, 2006; Cipriani et al., 2016; Friedrich, 2017; Turner et al., 2008; Zhou et al., 2015). By revealing that many different TSC-strength configurations, each associated with its own monoamine levels, could be equally well adapted to any chronic antidepressant treatment, the MS-model also identifies a key challenge to effective antidepressant drug design. By enumerating a very large set of possible neuroadaptive configurations to chronic drug or hormone administration, and by reporting the associated monoaminergic neurotransmitter and cortisol levels, our computational model can be used to address this challenge and to identify drug and/or hormone combinations that could be therapeutic for a higher proportion of patients, or patients with specific subtypes of depression requiring elevations in specific monoamines.

## 2. Materials and Methods

### 2.1 Model Formalism Overview

The MS-model takes the form of a recurrent network of units that all have the same sigmoidal (S-shaped, nonlinear but differentiable) activation function. This powerful computational formalism is a Turing-equivalent universal approximator that can be efficiently trained, via machine learning, to perform a broad range of desired, dynamic input-output transformations (Siegelmann and Sontag 1991). When used to model neural systems, the units in a recurrent network usually represent single neurons, or the average activity of the neurons in the same brain region (Anastasio, 2010). In modeling and analysis of the transmitter and hormonal systems that mediate mood, however, multiple levels of organization must be taken into account that include not only brain regions and neurons but also proteins and small molecules.

In the MS-model, individual units represent the brain regions, transmitters, receptors, transporters, enzymes, precursors, metabolites, and hormones that are involved in the pathologies of anxiety and depression. Some individual units represented whole brain regions that are central to these pathologies. The monoaminergic neurotransmitter-producing regions (DR, LC and VTA) were represented as single units because the majority of antidepressant drugs target these regions directly (Koenig and Thase, 2009). The HPA axis regions (PVN, pituitary gland, and adrenal gland) were represented as single units in order to incorporate the effects of the stress response in our analysis. The amygdala, prefrontal cortex (PFC), and hippocampus were represented because these regions are implicated in regulation of the HPA axis, and because they are also implicated in the antidepressant response itself through their involvement in cognitive control (Albert, Vahid-Ansari, & Luckhart, 2014; Dinan, 1996; Godlewska, Norbury, Selvaraj, Cowen, & Harmer, 2012; Malagi et al., 1996).

Single units also represented key neurotransmitters (e.g. 5HT, NE, and DA) and hormones (e.g. cortisol (CORT) and oxytocin (Oxt)), and many of the key transmitter receptors, transporters, and enzymes that are transmitter-system components (TSCs). See Supplemental Figure 1 for a diagram of the full MS-model.

Representing neurobiological entities (brain regions, transmitters, receptors, etc.) as single units enabled the model to represent the level (of activity, expression, concentration, etc.) of that entity in the whole brain, or in specific brain regions as appropriate. The 5HT unit, for example, represented the brain serotonin level, which is the major neurobiological endpoint for antidepressant action. The weights of the connections between the units also had specific identities in the model. For example, the effectiveness of the 5HT autoreceptor in inhibiting DR neurons was represented in the model by the absolute value of the inhibitory weight of the 5HT1A receptor (5HT1AR) unit onto the DR unit.

Relative to the overall number of network weights, a very small number of weights were of great significance because they represented TSCs whose efficacies, or strengths (expression levels, sensitivities, concentrations, synaptic locations, etc.), are known to adapt under chronic stress or chronic antidepressant administration. All of the weights in a trained network represent the normative strengths of influences of specific neurobiological entities on each other. The weights of the adaptable TSCs specifically were further adjusted to analyze the possible modes of adaptation to chronic stress, drugs, or hormones in the model.

### 2.2 Model Structure and Function

In describing neural networks it is necessary to distinguish inputs/outputs to/from the network overall, and inputs/outputs to/from individual units in the network. Units are categorized as input, output, or “hidden units.” There were 40 input units that provided input to the overall network. They take on assigned values and project to other units but do not receive connections from other units. Hidden and output units receive connections from input units and from each other. There were 23 output units that provided the output of the overall network. Output units are distinguished from hidden units in having targets (desired outputs). Hidden units have neither assigned nor desired values. The network consisted of 102 total units (input, hidden, and output).

Individually, the activity of each unit is a function of its inputs from the other units (this is not true for input units that do not receive inputs from other units). Each sending unit provides inputs to receiving units that are equal to the product of the sending unit’s activity level and the value of the weight (positive or negative) of the connection from the sending unit to the receiving unit. The net input to a receiving unit is the sum of the weighted inputs from all its sending units. The activity level of a unit is the value of its net input (i.e. weighted input sum) after it has passed through the sigmoidal activation function, which “squashes” the net input sigmoidally in the range between 0 and 1. The value of the sigmoidal squashing function for net input 0 is 0.50 (the midpoint of the 0-1 range).

The connection weights from the input, hidden, and output units to the hidden and output units are organized into the network weight matrix. The weight matrix consisted of 61 rows and 102 columns for a total of 6222 weights. Any input, hidden, or output unit could project to any hidden or output unit, but various distinctions between specific classes of connection weights were made in order to satisfy modeling goals.

The subject of the MS-model is the behavior of the monoaminergic transmitter systems and the stress hormone system (i.e. the monoamine-stress system) as they relate to the pathophysiology of depression. This behavior involves heavy interaction between the 3 monoaminergic transmitter systems and the stress hormone system, but also involves interactions between those systems and many other systems throughout the nervous system. The MS-model is therefore focused on the interactions between the monoaminergic transmitter and stress hormone systems, but also represents many other interactions that are included to ensure that model behavior is in line with a large set of empirical observations on the monoamine-stress system, as described in the literature.

In order to foreground monoamine-stress system interactions in the model, 3 classes of connection weights are distinguished: canonical, structure, and non-structure. Canonical weights are the weights of the known connections between units representing key components of the monoamine-stress system, such as the weight from the DR to 5HT, representing the effectiveness of the DR in producing 5HT. There were 23 canonical weights (see Supplemental Table 2). Structure weights are the weights of the connections between units representing neurobiological entities that are also known to interact empirically but are not canonical weights, such as the weight from DR to galanin, representing the effectiveness of the DR in co-releasing galanin. There were 305 total structure weights. Non-structure weights denote all other connection weights; they may or may not represent as yet unidentified interactions that actually do occur neurobiology. There were 5894 non-structure weights in total. The three classes of weights are treated differently during model training (see next subsection).

Two data structures were needed in order to construct and train the MS-model: a structure matrix and a truth table. Both the structure matrix and the truth table were compiled via a comprehensive literature search. The structure matrix is a two-dimensional matrix, coextensive with the network weight matrix, which specifies which neurobiological entities in the model are known to interact with which others, and their valance (positive or negative) if also known (see Supplementary Material S1: Details on Structure Connections). The structure matrix designates all structure and canonical weights in the model using non-zero integers. Non-structure weights take value 0 in the structure matrix. Figure 1 shows a highly simplified version of the structure of the model. The full model diagram can be viewed in Supplemental Figure 1: Complete Model Structure Diagram.

**Figure 1:**
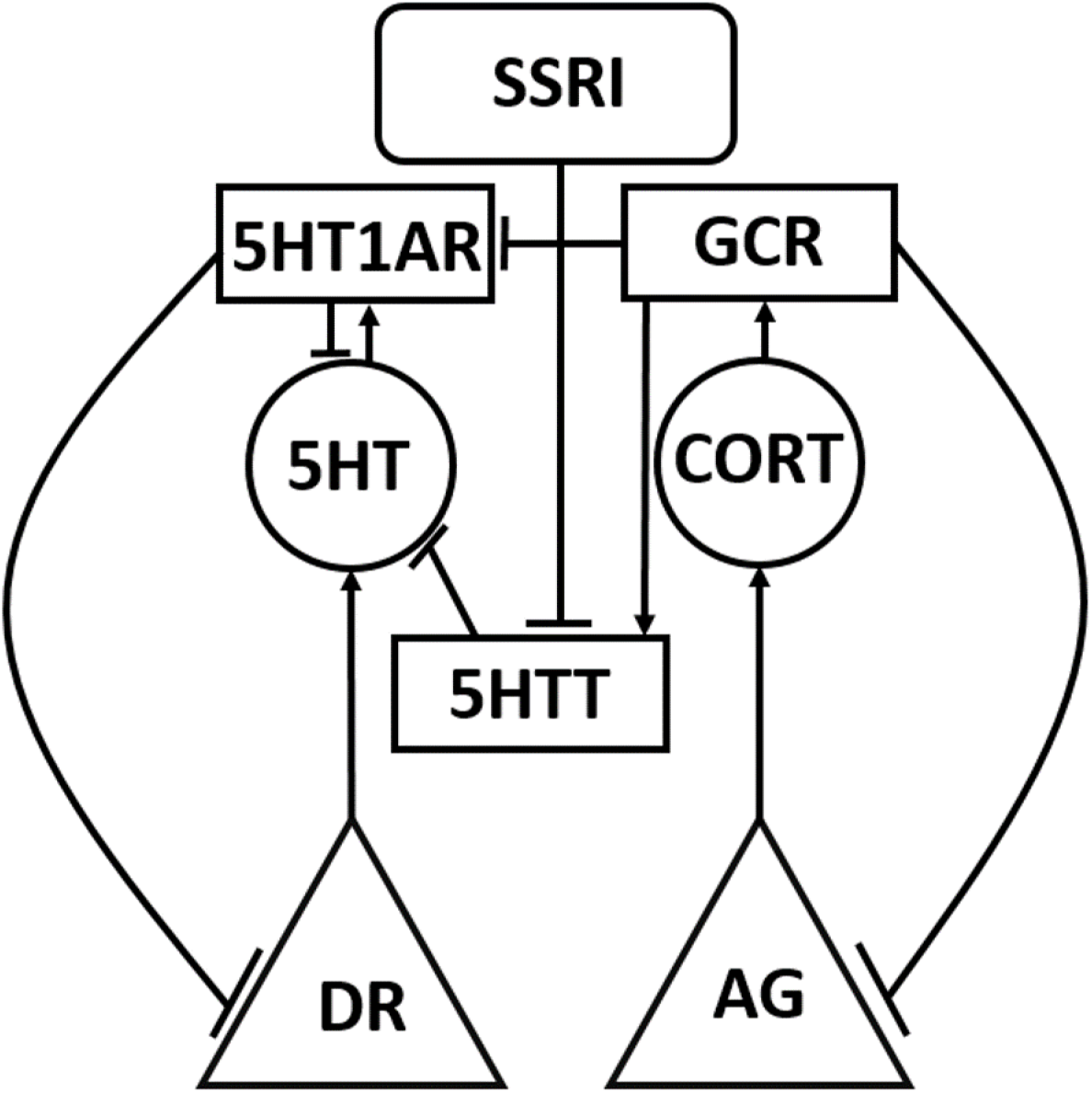
Simplified schematic representation of the model. The Monoamine-Stress model takes the form of a recurrent neural network of non-linear elements (units) that represent neurotransmitter-producing regions, enzymes, neurotransmitters, hormones, and receptors. Each unit type in the model is represented using a different shape in this highly simplified model diagram, in which only one or two of each unit type is shown. Neurotransmitter and hormone producing regions are represented as triangles, neurotransmitters and hormones are represented as circles, protein molecules are represented as rectangles, and inputs are represented as rounded rectangles. Connections between model units can be excitatory (arrow) or inhibitory (tee). Abbreviations: dorsal raphe, DR; adrenal gland, AG; serotonin, 5HT; cortisol, CORT; serotonin transporter, 5HTT; 5HT1A receptor, 5HT1AR; and glucocorticoid receptor, GCR.

The truth table is an array of input/desired-output training patterns that specifies how specific experimental manipulations, which are represented as patterns of network inputs, are known to affect specific neurobiological endpoints, which are represented as patterns of desired network outputs. Each row of the truth table represents the statistically significant results of one or more actual experiments. Inputs are either present or absent (1 or 0) and outputs are assigned discrete levels between 0.30 and 0.70. Outputs could either decrease maximally, decrease moderately, have no change, increase moderately, or increase maximally, corresponding to desired-output values of 0.30, 0.40, 0.50, 0.60, and 0.70, respectively (see Supplemental Material S2: Truth-table Justification for a summary of the experiments included in the truth table and corresponding references). Table 1 is a condensed, example input/desired-output table (i.e. truth table). The complete truth table has 66 input/desired output patterns (see the Excel file corresponding to Supplemental Table 1: Complete Model Truth-Table).

**Table 1:** Highly simplified example of the input/desired-output relationships used to train the model. The relationships represented in this truth table are based on the simplified diagram in Figure 1. Each row represents the consensus of the results of one or more experiments in which output levels were measured in response to each input. Row 1 is the baseline where there is no input. In rows 2 and 3 the inputs are drugs: SSRI or dexamethasone (glucocorticoid receptor agonist). In rows 4 and 5 the input is stress or adrenalectomy. The output values range from 0.30 to 0.70, where 0.30 represents maximal decrease, 0.40 represents moderate decrease, 0.50 represents the baseline value, 0.60 represents moderate increase, and 0.70 represents maximal increase.

| Row | Input | Desired output |  |  |  |
| --- | --- | --- | --- | --- | --- |
|  |  | DR | AG | 5HT | CORT |
| 1 | Baseline | 0.50 | 0.50 | 0.50 | 0.50 |
| 2 | SSRI | 0.40 |  | 0.60 | 0.70 |
| 3 | Dexamethasone |  |  | 0.60 | 0.30 |
| 4 | Stress | 0.60 | 0.70 | 0.60 | 0.70 |
| 5 | Adrenalectomy |  | 0.30 |  | 0.30 |

### 2.3 Training the Model

The model was trained using an efficient, gradient-based machine learning method known generally as recurrent back-propagation. The specific algorithm we used is due to Piñeda and it trains dynamic, recurrent neural networks to produce desired steady-state output patterns given specific input patterns (Pineda, 1987) (see Supplemental Material S3: Details on Model Training). The Piñeda algorithm assumes that networks reach steady-states and tends to train networks to reach steady states.

Prior to training, the complete network weight matrix, including canonical, structure, and non-structure weights, is randomized. Training occurs over 1×10^6^ training cycles (presentation of input/desired-output patterns). Input/desired-output patterns are presented to the network in random order during training. The steady-state network response to each network input is found after 100 iterations of unit updating (each unit computes the value of the sigmoid for its net input on each iteration). The differences between the desired and actual outputs of the units in the model were used to compute an error measure. The training method then computes the change for each weight using the error measure, and updates network weights on each training cycle.

Weight changes were scaled by a learning rate term before being applied to a weight. The learning rates were set to 1 for the canonical and structure weights, and to 0.10 for the non-structure weights, in order to disadvantage the non-structure connections because they are not known for certain to be involved in the behavior being modeled. All weights had an upper-bound at absolute value 10. All weights had a lower bound of 0 except for the canonical weights, which had a lower bound of 1 to ensure they exerted an influence on overall network performance. For more details on model training, see Supplemental Material S3: Details on Model Training.

We developed a pruning procedure to prune networks in order to eliminate unneeded non-structure connections. This was intended both to minimize the number of non-structure connections and to improve generalizability of model behavior. Generalizability was assessed by training the model on all single-manipulation inputs (e.g. single drugs) and testing on all combination inputs (e.g. drug combinations). The truth table included 41 single-input patterns and 25 combination-input patterns. Our pruning procedure eliminates 3981 non-structure connections (68% of all non-structure weights) on average and slightly improves generalization. The process of optimizing the pruning method is discussed in detail in Supplemental Material S4: Pruning Methods.

All further networks were trained using the following procedure: Networks with the full weight matrix (i.e. all canonical, structure, and non-structure connection weights) were trained on the full truth table (all single and combination inputs). Non-structure connections were pruned at the sensitivity cutoff optimized for generalizability, and the pruned networks were then retrained on the full truth table.

### 2.4. Adjusting Adaptable TSCs

Most antidepressants are administered chronically, so a model designed to represent the behavior of the monoamine-stress system in the context of depression must represent the responses to chronic-drug as well as to acute-drug administration. The MS-model can reproduce the effects of acute antidepressant administration by virtue of its training on the truth table, which includes many input/desired-output patterns that describe the effects of acute drugs. Because the monoamine-stress system is known to adapt under conditions of chronic drug (or chronic stress), the model must also account for neuroadaptation. Neuroadaptation is the process by which the overall activity levels (specifically the resting or spontaneous activity levels) of neurons in key brain structures are brought back to their normative baselines under conditions of chronic perturbation, such as chronic administration of a drug or other compound (see Introduction).

The 4 key brain regions in the MS-model are the 4 canonical neural structures: DR, LC, VTA, and PVN. These canonical brain structures are represented as individual units in the model. Neuroadaptation is known to occur through changes (i.e. adaptations) in the expression, sensitivity, efficacy, cellular or synaptic location, or other aspects associated with proteins (receptors, transporters, ion channels, etc.) that influence the activity levels of neurons in key brain regions (see Introduction). The 10 adjustable TSCs in the MS-model correspond to 10 actual TSC proteins that are known empirically to undergo adaptive changes under conditions of chronic manipulations within the purview of the model truth-table (see Supplemental Table 3: Adjustable TSCs). These 10 real TSCs are also known to play critical roles in the interactions between the 3 monoaminergic-transmitter systems and the stress-hormone system.

The strengths of cell-type specific TSCs are represented as individual weights in the MS-model. For example, the best known adjustable TSC in the context of antidepressant neurobiology is the 5HT1AR on DR neurons, which has been observed to “desensitize” (i.e. to become less sensitive to 5HT and therefore less effective in inhibiting the DR) (Blier and De Montigny, 1987; El Mansari et al., 2005). The TSC corresponding to the 5HT1AR on DR neurons is represented in the model as the weight of the connection from the 5HT1AR unit to the DR unit.

TSCs are network connection weights, but changes in these weights in the context of neuroadaptation are fundamentally different from changes due to neural network training. In neural network training, all connection weights can be changed in order to bring network outputs closer to specific desired, target outputs. In neuroadaptation, only a small subset of weights may change (i.e. those that correspond to key TSC proteins that are known to adapt), and these changes are directed not toward achievement of specific target outputs but to produce a more general restoration of the overall activity levels of neurons in key brain structures. For this reason, neuroadaptation was produced in the model simply by incrementally changing (i.e. adjusting up or down) the strengths of the network weights corresponding to the 10 adjustable TSCs. Note that the weights corresponding to the strengths of the 10 adjustable TSCs constitute a small number of the network connection weights.

An adjusted network was one in which the 10 weights corresponding to the adjustable TSCs are replaced with one or more adjusted values. Adjusted networks were considered adapted if their activation error was lower than their initial error, where initial error is the error due to acute administration of the drug or combination that is observed before any receptor adjustments have been made (see Results). The adjustment procedure was implemented “blindly” (i.e. in an unsupervised manner) so the adjustments could just as easily move the activities of the DR, LC, VTA, and PVN units away from normative activity as toward it, but adapted configurations were easily identified by the behavior of the adjusted networks.

We computed all possible configurations of the 10 adjustable TSC weights that were reachable by increasing or decreasing any single TSC weight by 0.50, within the predetermined TSC strength minimum of 0 and absolute maximum of 10, for a preset number of allowed adjustments that was the same for all weights. The “normative” strength of any network connection weight, including the weights corresponding to the 10 TSCs, is simply its value following neural network training. Due to the randomness inherent in neural network training (see previous subsection), equally well-trained networks can have very different network connection weights. This variability nicely corresponds to the natural variability in neurobiological properties that are known to occur between individuals (see Discussion).

To account for this inter-individual variability we studied adjustments of the normative weights in 3 representative networks, each trained from a different random initial weight matrix according to a different random order of input/desired-output presentations. We found all configurations of the 10 TSC weights reachable from each of the 3 representative, normative networks for all increments of 0.50 within the bounds of 0 and |10| up to a total of 6 adjustments, producing 382,747 total TSC-strength configurations over the 3 networks. Possible modes of adaptation of the MS-model were studied by analyzing this large set of adjusted configurations. Generation of the full set of TSC-weight configurations for a total of 7 adjustments, which would have produced over 11×10^6^ configurations, was not possible due to technical limitations (see subsection on Hardware Considerations).

### 2.5 Full-range Individual-weight Adjustment (FRIWA)

We assume that all normal individuals will adapt to chronic antidepressant but we further assume that different individuals will adapt in different ways, and not all of them will achieve “therapeutic” levels of the monoamines, defined as the levels necessary to achieve remission of depressive symptoms (see Discussion). It is also possible that some adjustable TSCs contribute more to the attainment of therapeutic monoamine levels than others. We define “therapeuticity” as the ability of a TSC to contribute to the attainment of therapeutic monoamine levels through adjustments in its strength. We conducted full-range individual-weight adjustment (FRIWA) analysis (a kind of sensitivity analysis) to gauge the therapeuticity of single TSCs.

The starting configurations for FRIWA analysis were all of the configurations adapted to chronic SSRI that were also therapeutic, extracted from the exhaustive set of configurations of adjustments in all TSCs up to a total of 6 adjustments. The total number of adapted and therapeutic configurations over all 3 representative networks was 5598. Therapeutic configurations were those that increased 5HT above the 5HT therapeutic floor (>=0.70) and decreased CORT below the therapeutic CORT ceiling (<=0.70).

FRIWA analysis involved adjustment of the weight of a single adaptable TSC across its full range (0 to |10|; note that individual TSCs are either positive or negative), while the weights for the 9 other adaptable TSCs remained frozen at their starting values, in each of the 5598 adapted and therapeutic starting configurations. FRIWA occurred in steps, where each individual weight adjustment (IWA) was an increment of |0.50|. Thus, FRIWA generated a set of 200 new configurations (20 IWA adjustments for each of 10 TSC weights) starting from each of the 5598 adapted and therapeutic configurations for a total of 980,193 new configurations. All configurations that were no longer adapted after a step of IWA were excluded from further analysis, leaving 493,564 adapted TSC-strength configurations. Of those, 285,635 configurations were designated “resistant,” because they remained therapeutic despite a step of IWA, while the remaining 207,929 configurations were designated “sensitive” because there were rendered non-therapeutic by a step of IWA.

The weights of each of the 10 adjustable TSCs in all of the post-IWA configurations were pooled over each representative network, and separated on the basis of resistance and sensitivity, excluding the weights that were manipulated by FRIWA. The mean weight of each adjustable TSC was computed for both the resistant and sensitive configurations of each network and compared (see Results). Further FRIWA analysis involved computing the pairwise correlations between all TSC weights over all the resistant, or over all the sensitive, configurations in each network. Only pairwise correlations that were statistically significant (p < 0.05) over all 3 representative networks would have been reported. Again for correlation analysis, the TSC weights that were manipulated by FRIWA were excluded.

### 2.6 Temporal-logic Model-checking

As described above, we studied the consequences of adaptation by generating a large set of adapted TSC-strength configurations (see subsection on Adjusting Adaptable TSCs). We also studied the process of adaptation by making allowed TSC weight adjustments (increments of 0.50 up or down, within bounds of 0 and |10|) in all possible sequences. Linear temporal logic (LTL) is a type of modal temporal logic which allows for reasoning about sequences of discrete states evolving in time. LTL analysis allows the evaluation of temporally specified logical propositions such as whether a specific state is always maintained or eventually reached; whether a specific state pertains only until another state pertains; or whether a specific state always leads to another specific state. Temporal-logic model-checking allows us to determine such temporal relationships between TSC-strength configurations (i.e. states).

We used LTL model checking to determine if specific degrees of neuroadaptation (e.g. a configuration in which a specific TSC has been adjusted down 3 times) always leads to an adapted and therapeutic configuration (for all possible sequences of adjustments proceeding from that configuration up to a total of 6 adjustments) (see Supplemental Material S5: Details on Temporal-logic Model-checking Procedure for details on model-checking statements).

In order to evaluate model-checks in the MS-model, it was necessary to define a set of logical predicates to be tested. The following predicates were used in the temporal-logic analysis: `fht_high`, 5HT is above the 5HT therapeutic floor (>=0.70); `cort_low`, CORT is below the therapeutic CORT ceiling (<=0.70); `TSC_sens_gt_3`, the adjustable TSC has sensitized by at least 3 steps; and `TSC_desens_gt_3`, the adjustable `TSC` has desensitized by at least 3 steps. Note that TSC in the last two predicates is a placeholder for any specific, adjustable TSC (e.g. 5HT1AR).

Then we evaluated the following propositions for each of the 10 adjustable TSCs, where `| ->` and `/\` are the LTL “LEADS TO” and “AND” operators, respectively:

~~~
TSC_sens_gt_3 |-> fht_high /\ cort_low
TSC_desens_gt_3 |-> fht_high /\ cort_low
~~~

These are equivalent to the LTL propositions that if at any point in the trajectory of TSC strength adjustments, the TSC sensitizes or desensitizes by at least 3 increments, then 5HT is high and CORT is low at some subsequent point in time. We found that both these propositions were false for all of the tested TSCs in all 3 representative networks (see Results).

Next, we evaluated the following more-easily-satisfiable propositions, where `\/` and `~` are the LTL “OR” and “NOT” operators, respectively:

~~~
TSC_sens_gt_3 |-> (fht_high /\ cort_low) \/ ~TSC_sens_gt_3
TSC_desens_gt_3 |-> (fht_high /\ cort_low) \/ ~TSC_desens_gt_3
~~~

These are equivalent to the LTL propositions that if at any point in the trajectory of TSC strength adjustments, the TSC sensitizes or desensitizes by at least 3 adjustments, then either 5HT is high and CORT is low at some subsequent point in time, or the TSC is no longer sensitized or desensitized. If 1 adjustable TSC was solely responsible for modulating the 5HT and CORT levels, then the model check for either of those 2 propositions for that specific, adjustable TSC should return True. Each LTL model check was carried out using each of the 3 representative networks for up to 6 adjustments, and only model-checking results that were consistent over all 3 networks are reported in Results.

### 2.7 Hardware Considerations

All MATLAB computational procedures were performed on an Intel Core 2 Duo CPU processor with 2, 2.33 GHz cores and 4.00 GB of RAM under the Windows 7 operating system, an Intel-inside CORE i7 processor with 2, 2.69 GHz cores and 8.00 GB of RAM under the Windows 8 operating system, and an Intel Core i7 processor with 4, 4.00 GHz cores and 32.00 GB of RAM under the Windows 10 operating system. MATLAB was used for training, pruning, enumerating, and analyzing all adjustable-TSC configurations with up to 6 adjustments in TSC strength, generation of all histograms and heatmaps, and FRIWA analysis. Training a network (100 time steps, 1×10^6^ iterations, 66 training patterns) took between 20 and 25 minutes. Exhaustively computing full sets of adjustable TSC configurations to degree 6 took 1 minute. Computational overhead (memory limitations) prevented computing the full set to degree 7. Checking the set of TSC-strength configurations for neuroadaptation to chronic drug or hormone combinations in each of the 3 representative networks took 8 minutes per network using MATLAB.

Python was used for enumeration of TSC-strength configurations and LTL analysis. Python search and LTL analysis of TSC-strength configurations having 127,582 states took 34 seconds on one quad-core Intel Core i5 processor. For subsequent model checks where steady-state values had been cached, model checks took an average of 3.10 seconds. Python enumeration of TSC-strength configurations was limited to degree 6 for consistency with the MATLAB analysis and because the results were unlikely to be different for degree 7 than for degree 6 (see Results).

## 3. Results

The results are based on analysis of a large set of configurations of the strengths of specific transmitter system components (TSCs), which are represented as connection weights in the monoamine-stress network model (MS-model; see Methods for details). The main modeling assumption is that the real monoamine-stress system will adapt to chronic conditions (chronic stress, chronic drug administration, etc.) by adjusting the strengths (through changes in expression, sensitivity, synaptic location, etc.) of TSCs in order to move brain-region activity levels closer to their normative (no-stress, no-drug, etc.), baseline levels. To avoid making unjustifiable assumptions, the strategy pursued here was to make all possible TSC adjustments in order to make all possible configurations of TSC strengths, given certain practical constraints on the number of configurations. TSC-strength configurations that conferred specific properties on the MS-model were then identified. The 5 properties of interest in the MS-model were the level of adaptation, the levels of the 3 monoaminergic transmitters, and the level of CORT as represented in the model.

TSC-strength configurations were either analyzed in groups, in order to determine the distributions of configurations that exhibited specific properties, or in sequences of TSC-strength adjustments, in order to analyze possible sequential (temporal) relationships between TSC-strength configurations. The adjustable TSCs are a set of 10 TSCs that are known to adjust empirically under the conditions of the experiments from which the model training patterns were derived. As we found (see below in this section), these 10 TSCs largely overdetermined the 5 properties of interest in the MS-model.

To account for the possibility that different initial configurations of the weights that represent TSC strength in the model (and of non-TSC weights also) could affect the results, we analyzed 3 networks, all trained to similarly high degrees of accuracy in reproducing desired behavior but from different initial random weight matrices and according to different random orders of training pattern presentations. The model illustrates how clinically observed heterogeneity in the antidepressant response could reflect heterogeneity in adapted TSC-strength configurations that are associated with widely divergent monoamine levels.

The model provides predictions concerning the efficacy of specific drug or drug and hormone combinations that could potentially be more therapeutic than single antidepressants. It also provides insight into the chronic effects of drug or drug and hormone combinations on the levels of different monoamines to aid in designing more targeted antidepressant therapies for different depressive subtypes. Results also provide insight into the therapeuticity (i.e. the ability to contribute to a therapeutic state) of different TSC types, through analysis of large sets of adapted configurations and temporal relationships between configurations that occur during the process of neuroadaptation.

### 3.1 Agreement between Actual and Desired Outputs

Networks were trained, pruned, and then retrained as described in Methods. The results of training can be viewed in Figure 2. Each plot in Figure 2 shows all of the desired and actual outputs for one brain-region, transmitter, or hormone output unit (DR, LC, VTA, PVN, 5HT, NE, DA, or CORT). Each actual output response and desired output response is plotted as a solid line and a dashed line, respectively. Correspondence between steady-state actual responses and desired responses is nearly exact. The average error of these units over all of the trained-pruned-retrained networks was very low (6.39×10^−5^). The heatmaps in Supplementary Figure 2 illustrate the extent of model agreement for all output units over the whole training set both before (Supplementary Figure 2A-C) and after pruning (Supplementary Figure 2D-F).

**Figure 2:**
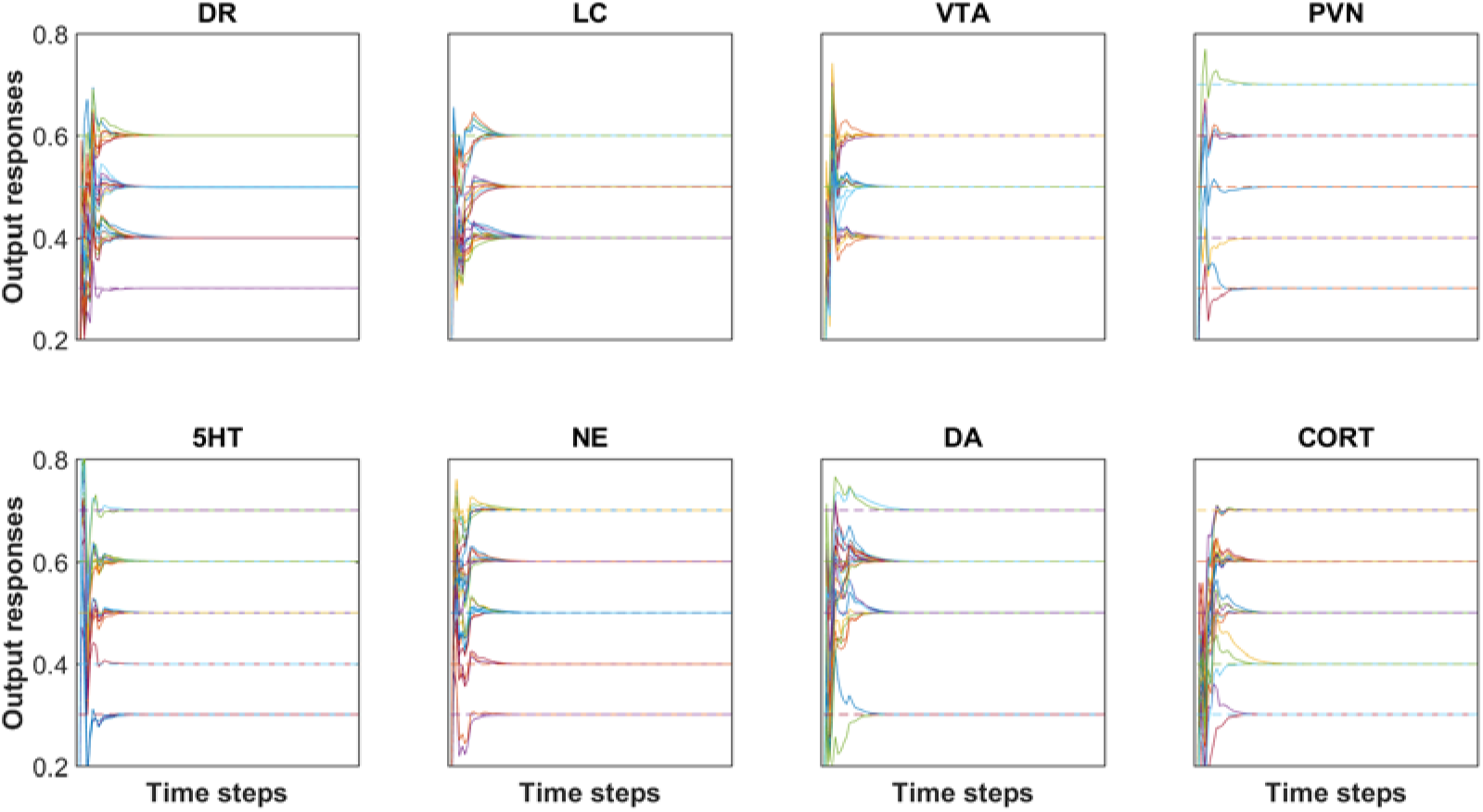
Agreement between desired (i.e., target) and actual outputs after pruning and re-training. All of the desired and actual outputs, collected over all input/desired-output patterns, are represented in a single plot for each of the brain region and transmitter or hormone output units (DR, LC, VTA, PVN, 5HT, NE, DA, and CORT). Each output response is plotted as a solid line and each target (i.e. desired) output is plotted as a dashed line. Each of the outputs reach a steady-state value within 25 time steps. The RMS error of this network is 5.10×10^−5^. Note that the solid line (actual output) is superimposed on the dashed line (desired or target output), illustrating the accuracy of the training method.

### 3.2 Enumeration of Adjustable TSC-strength Configurations

The baseline activity levels of key model units are represented as dashed blue lines in Figure 3. The baseline activity is simply the activity of units in a trained network when the inputs to the network are all 0. All units have positive (nonzero) baseline activities because the squashing function, which determines each unit’s output as a function of its net weighted input, produces a “spontaneous” output activity of 0.50 for a net input of 0. Note that the squashing function bounds unit activation between 0 and 1 (see Methods).

**Figure 3:**
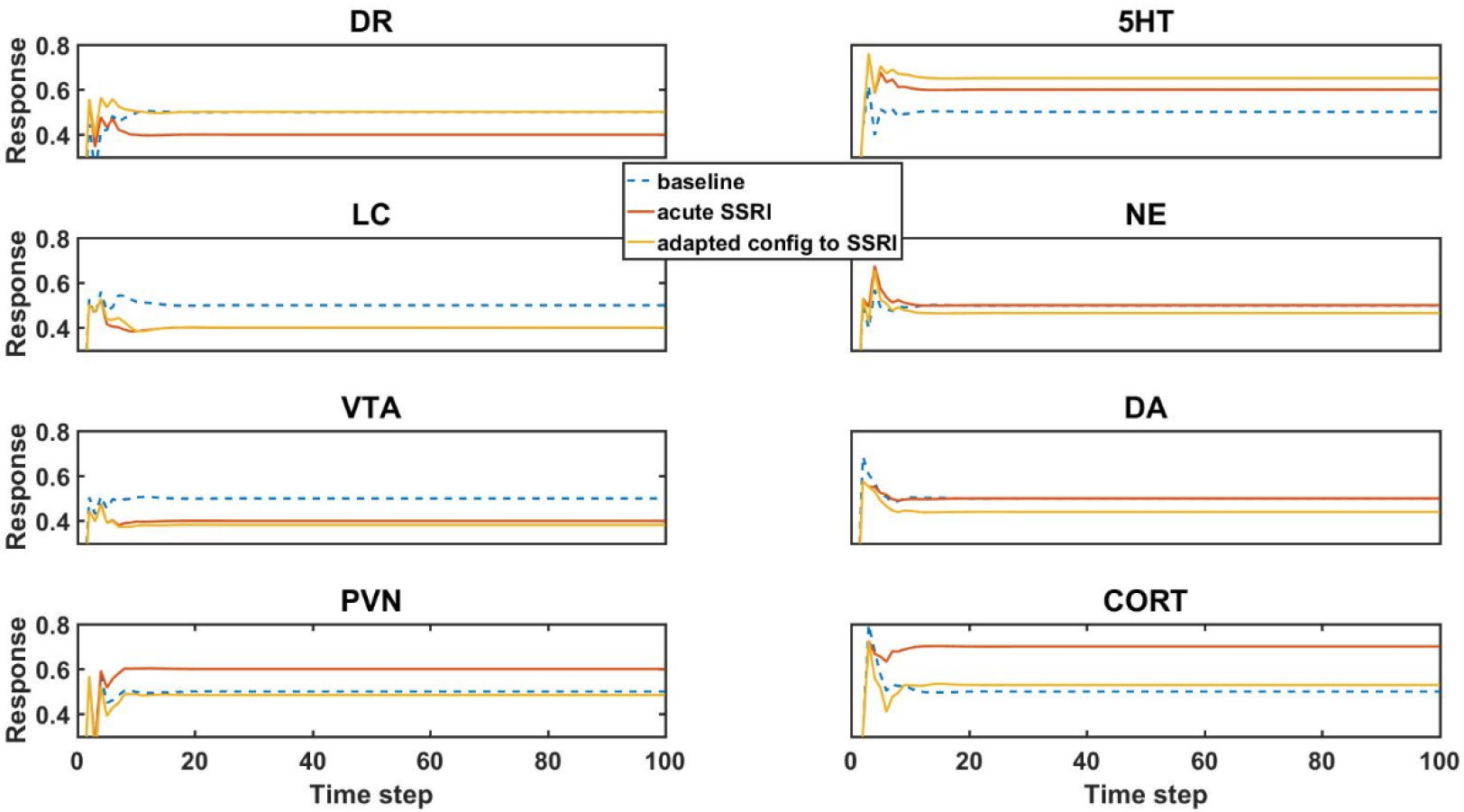
Model element activities in the baseline (no-drug) condition, acute (no-adaptation) SSRI condition, and chronic (adaptation) SSRI condition. Each plot corresponds to a different model unit as labeled. The blue dotted line in each plot shows the baseline activity level of a unit in the normal (no-drug) baseline condition. The red line in each plot shows the unit activity in the acute (no-adaptation) SSRI condition. Note that acute SSRI changes the activity levels of all of the units. The yellow line in each plot shows the adapted activity of a unit in an example, adapted configuration with chronic SSRI. Note that the adapted DR and PVN unit responses return closer to baseline, and the adapted 5HT and CORT responses increase and decrease, respectively. Abbreviations: dorsal raphe, DR; locus coeruleus, LC; ventral tegmental area, VTA; paraventricular nucleus of the hypothalamus, PVN; serotonin, 5HT; norepinephrine, NE; dopamine, DA; cortisol, CORT.

The units in the network influence each other’s activity through their weighted interconnections. Because the unpruned weights of the connections between the units are nonzero (positive or negative), the baseline activities of the units can depart substantially from 0.50. The baseline activity of the units is basically the “response” of the network to 0 input. Following an initial transient, all unit “responses” to 0 input (baseline responses) settle into a stable activity pattern within 25 time steps that is maintained for the duration of the response. The baseline activity levels of the units in a trained network are their “normative” activity levels.

Simulated acute administration of an SSRI (an SSRI input level of 1) alters the responses of the units in the model, represented as orange lines in Figure 3. Note that unit activities can deviate substantially from their normative baselines due to acute SSRI. From the neuroadaptive standpoint, any deviation from normative baseline is considered an error that should be corrected by the adaptive process. We define “adaptation error” as the sum of the absolute differences from baseline of the activities of the DR, LC, VTA, and PVN units, because these are the canonical units in the MS-model.

Chronic (as opposed to acute) administration of a drug (or combination) was simulated simply by keeping the input unit corresponding to that drug (or drugs, for a combination) at 1. We further defined “initial error” as the adaptation error of a trained network subjected to chronic administration of a drug (or combination) in the absence of any adjustments of the strengths of the weights representing the adjustable TSCs. An “adapted network” was any network that, due to one or more adjustments in TSC weights, had adaptation error lower than initial error. Model responses to acute SSRI, and adaptation to chronic SSRI, were of particular interest because SSRI is the drug class most often prescribed for the treatment of anxiety and depression. The responses of the canonical units in an example network adapted to chronic SSRI are represented in Figure 3 as yellow lines. This figure shows that adaptation to an SSRI *can* return the responses of DR and PVN back toward normal while also increasing 5HT levels and decreasing CORT levels, which is in agreement with clinically observed chronic SSRI effects (Ceglia et al., 2004; Nikisch et al., 2005; Rush et al., 2006; Rutter et al., 1994).

Many other adapted networks, however, did not also have this pattern of adapted behavior. Transmitter and hormone responses to chronic drug administration, therefore, must be evaluated in many different TSC-strength configurations, derived from adjustments in the TSC weights of several different initial networks. As described above, our results are based on 3 representative networks (see also Methods).

We analyzed all of the configurations, starting from each of the 3 different, representative networks that could be generated by up to 6 adjustments at an increment of 0.50 (the number of adjustments was limited to 6 due to computational overhead, see Methods). The distributions of the 3 monoaminergic transmitter and CORT levels corresponding to all the adapted configurations of the 3 networks to chronic SSRI are shown as histograms in Figure 4. Each histogram in Figure 4 shows the numbers of configurations adapted to chronic SSRI that had levels of 5HT, NE, DA, and CORT falling within various bins as indicated. The first 3 columns show the histograms for the 3 representative networks separately, while the fourth column shows the histograms for the 3 representative networks combined. In this histogram figure and in all subsequent histogram figures the baseline, average, and therapeutic levels of each transmitter or hormone will be represented with a blue, green, or magenta line, respectively.

**Figure 4:**
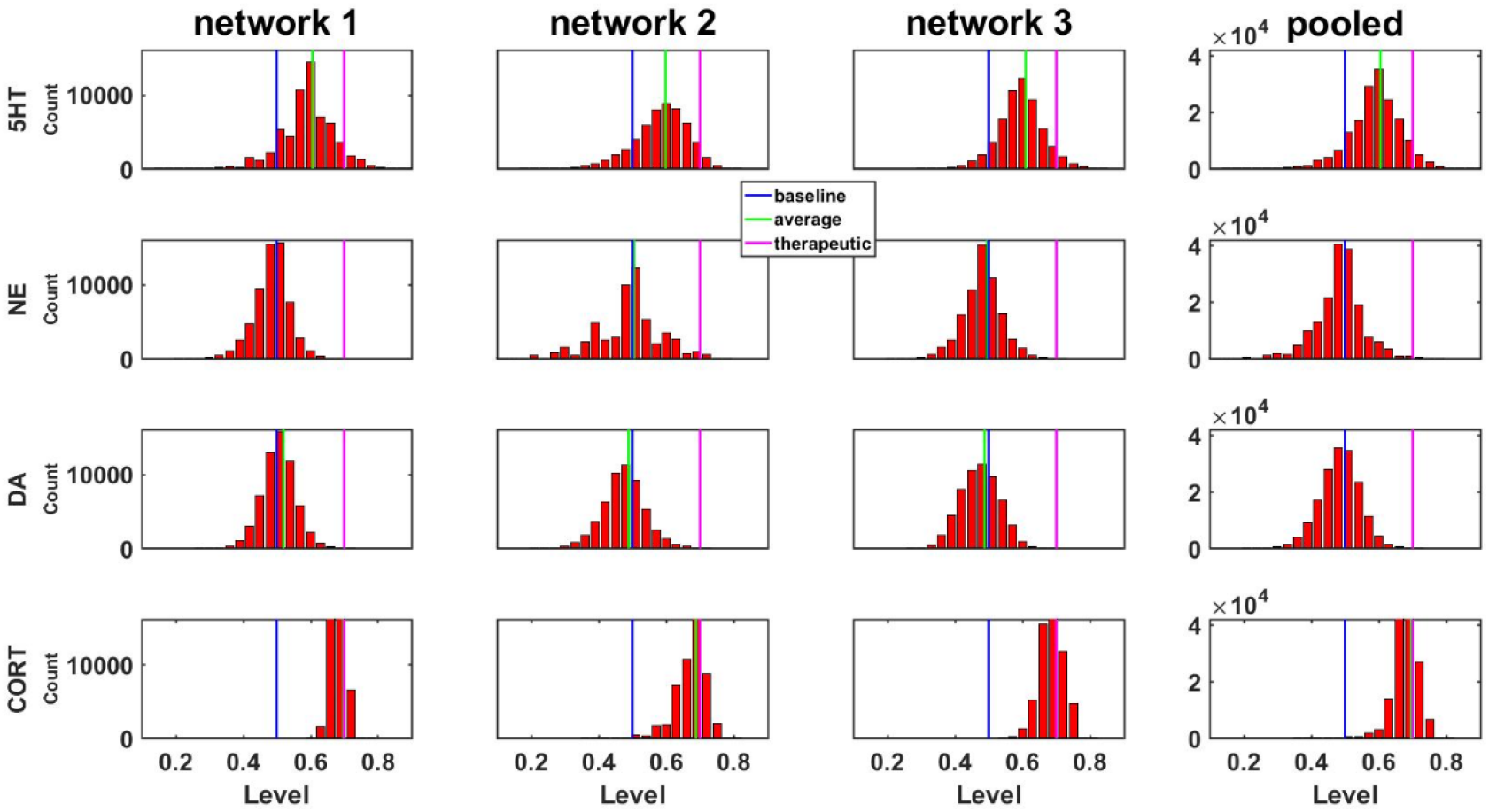
Histograms showing numbers of configurations adapted to SSRI expressing different levels of monoamines or CORT. Results are shown for 3 networks individually or pooled. Networks were adapted to chronic SSRI. The bin width of these and all other histograms was set to 0.03. The blue, green, or magenta vertical line in each plot is located at the baseline level (0.50), the average level, or the therapeutic cutoff (0.70), respectively, for each neurotransmitter or hormone. This figure also demonstrates pooling of the adapted configurations from 3 networks.

Although “therapeutic” monoamine levels have not been determined, it has been observed that 5HT levels with chronic SSRI are about double the level with acute SSRI (Ceglia et al., 2004). Because the desired baseline for 5HT was set to 0.50, and the desired 5HT output for acute SSRI was set to 0.60 (see Supplemental Material S2: Truth Table Justification), the therapeutic 5HT floor was set to 0.70 in order to double the acute increase of 0.10. For consistency, 0.70 was also used as the therapeutic floor for NE and DA because therapeutic levels for NE and DA with chronic antidepressant have not been identified conclusively. CORT levels have been found to decrease with chronic SSRI administration, so the therapeutic ceiling for CORT was set to 0.70 to reflect a decrease from the acute-SSRI CORT level of 0.70 (Lenze et al., 2011; Ruhé et al., 2015).

A central goal of this study was to use the trained network models to evaluate the possibility that combinations of antidepressant drugs, or combinations of drugs and hormones, could be more therapeutically effective that single SSRIs. Configurations adapted to chronic administration of select drugs and hormones in combination with SSRI were analyzed. The results of some specific drug and hormone combinations are shown in Figures 5 and 6. The rows of these 2 sets of histograms (Figures 5 and 6) show the monoamine and CORT distributions of all configurations starting from all 3 representative networks that were adapted to SSRI alone, to SSRI paired with another drug or hormone, and SSRI combined with the two other drugs or a drug and a hormone. Each column of these 2 sets of histograms (Figures 5 and 6) corresponds to a specific transmitter or hormone unit (5HT, NE, DA or CORT). Viewing the histograms down the columns shows how the distributions of monoamine and CORT levels are affected by the different drugs and combinations.

**Figure 5:**
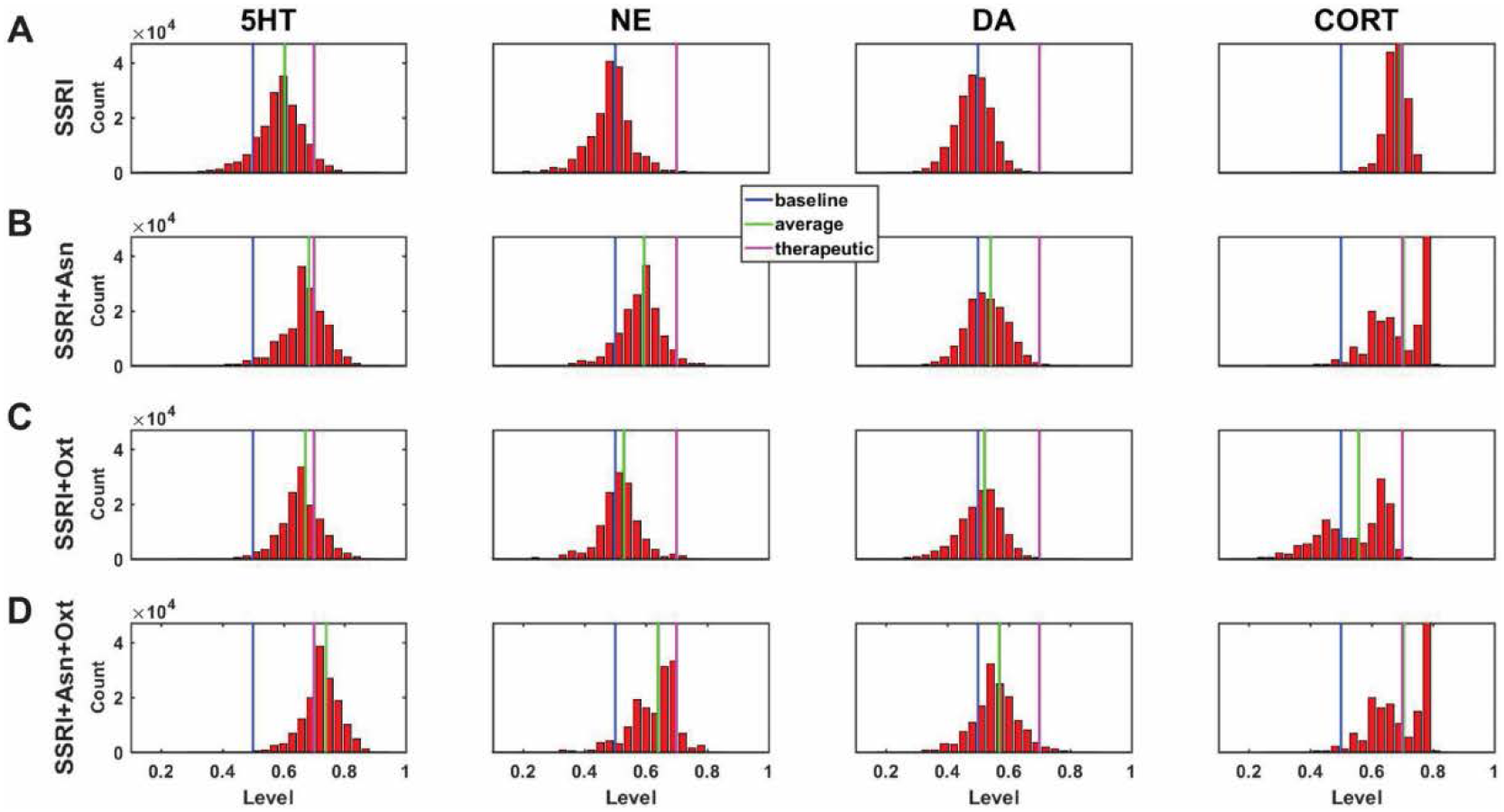
Histograms showing number of adapted configurations expressing different monoamine and CORT levels with combinations of SSRI, Asenapine, and Oxytocin. Networks were adapted to SSRI alone (A), SSRI+Asenapine (Asn, an antipsychotic drug) (B), SSRI+Oxytocin (Oxt, a hormone) (C), and SSRI+Asn+Oxt (D). (B) and (C) show that combining an SSRI with either Asn or Oxt increases the proportion of high monoamine and low CORT states over the SSRI by itself. (D) shows that combining an SSRI with both Oxt and Asn further increases the proportion of high monoamine and low CORT states. These histograms suggest that combining an SSRI with either Oxt, Asn, or both may be therapeutic for a greater proportion of patients than an SSRI administered alone.

Figure 5A shows the results of adaptation to SSRI alone, 5B shows SSRI paired with Asenapine (an antipsychotic drug), 5C shows SSRI paired with Oxytocin (a peptide neurohormone), and 5D shows SSRI combined with both Asenapine and Oxytocin. Figure 5B shows that combining SSRI with Asenapine not only increases the proportion of adapted states with therapeutically elevated (toward the right) 5HT, but also shifts the NE histogram to the right as well increasing the proportion of adapted states with elevated NE. Figure 5B also shows that the combination of SSRI and Asenapine can decrease CORT levels in a higher proportion of adapted configurations below the therapeutic ceiling (toward the left). In Figure 5C, the combination of an SSRI and Oxytocin also shifts the 5HT distribution to the right and increases the proportion of adapted states with low CORT. The combination of all 3 factors (SSRI, Asenapine, and Oxytocin) in Figure 5D increases the proportion of adapted states with high monoamine levels for all 3 monoamines, and also increases the proportion of adapted states with low CORT beyond the levels observed with SSRI by itself.

Figure 6A shows the results of adaptation to SSRI alone, 6B shows SSRI paired with Bupropion (an atypical antidepressant), 6C shows SSRI paired with Olanzapine (an antipsychotic drug), and 6D shows SSRI combined with both Bupropion and Olanzapine. Figure 6B shows that adaptation to the combination of SSRI and Bupropion can increase the proportion of adapted states with high NE and DA levels over that observed with the SSRI by itself. It also shows that this combination decreases CORT levels in a higher proportion of adapted states than the SSRI by itself. Figure 6C shows that the combination of SSRI and Olanzapine can increase the proportion of adapted states with therapeutic 5HT over the SSRI by itself. The combination of all 3 (SSRI, Bupropion, and Olanzapine) in Figure 6D was found to shift all 3 of the monoamine (5HT, NE and DA) histograms to the right (toward high monoamine levels) and increase the proportion of adapted states that reduce CORT below its therapeutic ceiling. Overall, these histograms illustrate how combinations of chronic drugs (or of drugs and hormones) can increase the proportion of adapted states with elevated monoamines and reduced CORT levels (see Discussion).

**Figure 6:**
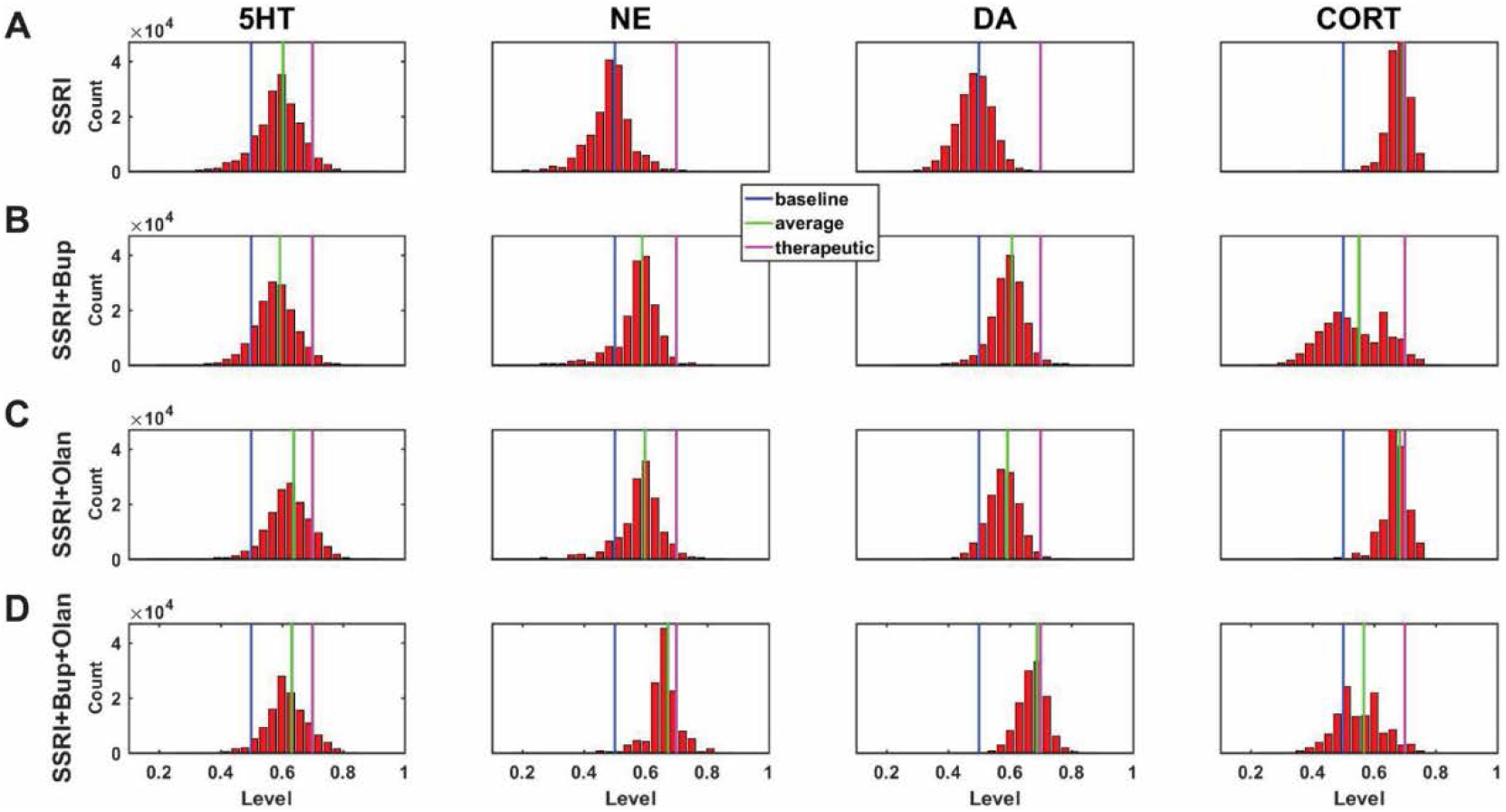
Histograms showing number of adapted configurations expressing different monoamine and CORT levels with combinations of SSRI, Bupropion, and Olanzapine. Networks were adapted to SSRI alone (A), SSRI+Bupropion (Bup, a NET and DAT blocker) (B), SSRI+Olanzapine (Olan, an antipsychotic drug) (C), and SSRI+Bup+Olan (D). (B) and (C) show that combining an SSRI with either Bup or Olan increases the proportion of high monoamine and low CORT states over the SSRI by itself. (D) shows that combining an SSRI with both Bup and Olan further increases the proportion of high monoamine and low CORT states. The histograms in (D) illustrate that the combination of SSRI+Bup+Olan shifts the monoamine and CORT histograms, especially those of NE and DA, toward more therapeutic states, suggesting that this combination can be therapeutic for a greater proportion of patients than an SSRI administered alone.

### 3.3 Predicting the Efficacy of Drug and Hormone Combinations

The model was then used to evaluate the configurations adapted to chronic administration of 60 drug or drug/hormone pairs and triples. The 27 pairs consisted of SSRI paired with each of the 27 drugs and hormones that were used as individual (single) inputs in model training. The 33 triples consisted of the subset of those pairs that have been used to augment SSRI action clinically or experimentally (in 1 case), combined with a third substance that was either Oxytocin (a hormone), Antalarmin (a CRF1 receptor antagonist), or Olanzapine (an antipsychotic). These 3 substances were selected due to recent preliminary evidence suggesting that Oxytocin, Antalarmin, and Olanzapine could potentially have antidepressant effects either by themselves or in combination with another antidepressant (Amini-Khoei et al., 2017; Björkholm et al., 2015; Scantamburlo et al., 2015; Thomas et al., 2017).

The monoamine levels for the 3 networks adapted to each of the drug or hormone pairs and triples were compiled and the average monoamine levels were computed. The average levels of each of the 3 monoamines for each chronic drug or hormone combination are shown as rows in the heatmap in Figure 7. Each column represents the level of one monoamine (5HT, NE, or DA, moving across the columns). For purposes of illustration, quantification, and ordering, the 3 monoamine levels can be combined into a monoamine vector: [5HT NE DA]. Baseline, therapeutic, and excess monoamine reference vectors can then be defined. The baseline reference vector consists of the baseline monoamine levels [0.50 0.50 0.50], the therapeutic reference vector consists of the therapeutic monoamine levels [0.70 0.70 0.70], and the excess monoamine reference vector consists of excessively high monoamine levels [0.80 0.80 0.80]. The excess monoamine reference vector represents a tripling of the acute level and is included in order to identify drug pairs and triples that may elevate the monoamines high enough to produce unwanted side effects (Boyer and Shannon, 2005; Shrier et al., 2000).

**Figure 7:**
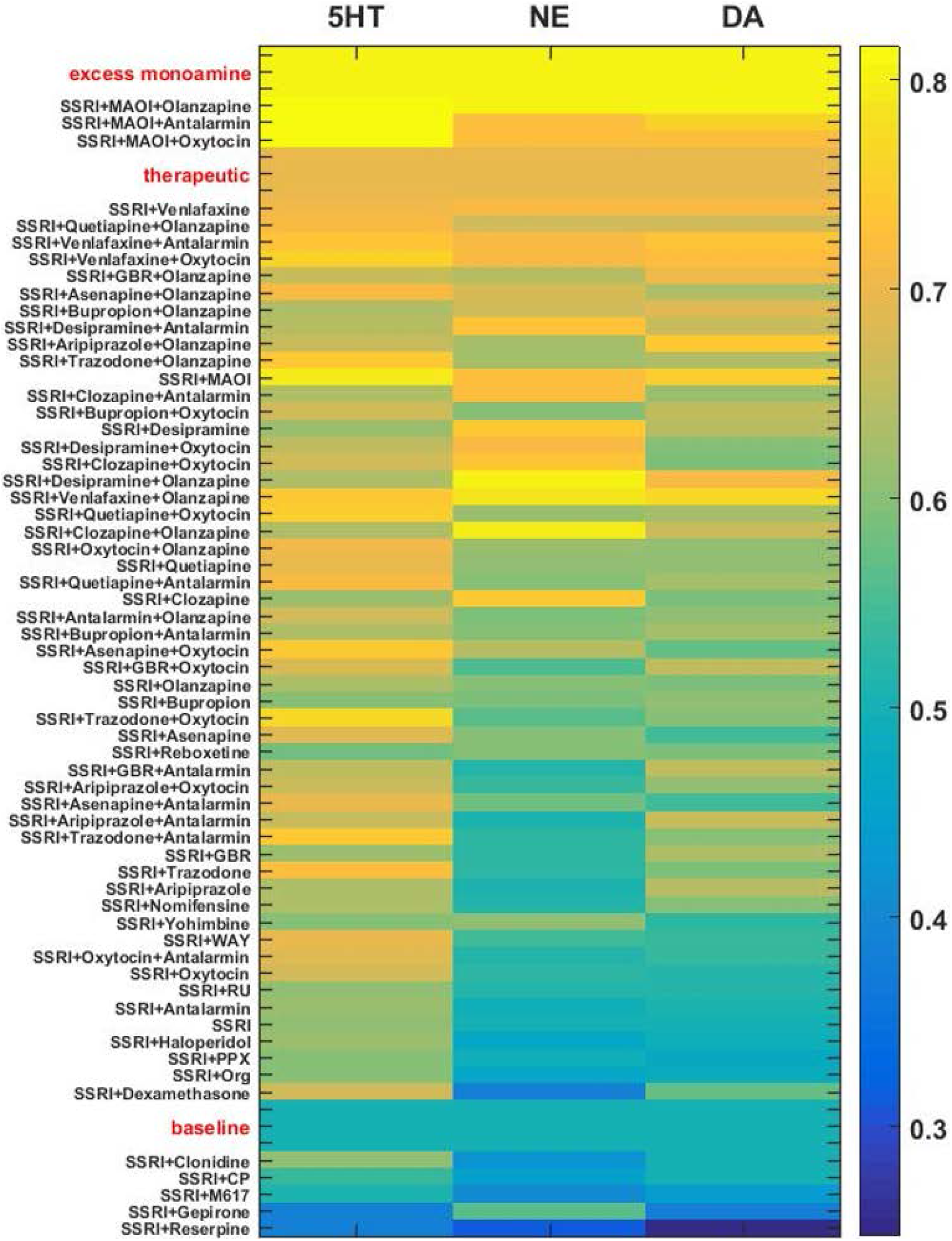
Heatmap of adapted monoamine levels with SSRI, all other drugs paired with SSRI, and selected 3-drug combinations. Adapted monoamine levels were averaged over the 3 networks and expressed as a vector [5HT NE DA]. The excess monoamine reference vector, representing levels high enough that they could be associated with unwanted side effects, was set to [0.80 0.80 0.80]. All drug combinations that resulted in 1 or more excess monoamine levels were ordered by vector distance from the excess monoamine reference vector. The therapeutic and the baseline reference vectors were set to [0.70 0.70 0.70] and [0.50 0.50 0.50], respectively. The baseline reference vector and all remaining drug pair and triple vectors were ordered by vector distance from the therapeutic reference vector. Abbreviations: GBR-12909, GBR; WAY-100635, WAY; Pramipexole, PPX; RU-28362, RU; Org-34850, Org; CP-96345, CP; Monoamine oxidase inhibitor, MAOI.

All adapted monoamine vectors [5HT NE DA] that had any monoamine elevated above 0.80 were ordered by their vector distance from the excess monoamine vector; all remaining adapted monoamine vectors were ordered by their vector distance from the therapeutic reference vector. This figure can be used as a tool to help predict monoamine levels following adaptation to chronic administration of each of the selected drug combinations, and to identify potentially effective interventions for the different subtypes of depression that are believed to respond to elevations in different monoamines (Malhi et al., 2005) (see Discussion).

### 3.4 Full-range Individual-weight Adjustment (FRIWA) to Evaluate the Therapeuticity of Each Adjustable Weight

Though we identified and examined adjustments in 10 adjustable TSCs, it is possible that single TSCs can alone determine the therapeutic state of the MS-model, regardless of the values of the weights representing the other adaptable TSCs. We define “therapeuticity” generally as the ability of a biological factor to alter the properties of a biological system in a therapeutic direction. Specifically here, therapeuticity is the ability of a TSC to alter 3 of the properties of interest in the MS-model: the level of adaptation, the level of 5HT, and the level of CORT. Identification of the TSCs that could alone determine therapeutic state could enhance antidepressant drug design by identifying specific receptors or transporters that could be targeted in single-drug therapies.

In order to determine if therapeutic effects were specifically dependent on single adjustable TSCs, an analysis was conducted that evaluated the contribution of each individual, adjustable TSC to therapeutic adaptation with chronic SSRI administration. Therapeutic states were defined as adapted states that increased 5HT up to or above the therapeutic 5HT floor (>=0.70) and decreased CORT down to or below the therapeutic CORT ceiling (<=0.70). All configurations, generated from adjustments of all adjustable TSC weights from 1 to 6 adjustments, which were adapted to chronic SSRI and also therapeutic were pooled for each of the 3 representative networks for full-range individual-weight adjustment (FRIWA) analysis.

In FRIWA analysis, for each adapted and therapeutic configuration, 1 adjustable TSC weight was adjusted across its full range (0 to absolute value 10), while the 9 other TSC weights remained frozen. FRIWA occurred in 20 individual weight adjustment (IWA) steps of 0.50 each (see Methods). All states that were no longer adapted after an IWA step were excluded from the analysis. The still-adapted configurations that were resistant (remained therapeutic after an IWA step) or sensitive (became non-therapeutic after an IWA step) were then identified. The weights for each TSC were compiled for all resistant or all sensitive post-FRIWA configurations for each representative network. The average values of the weights for each TSC over either the resistant or the sensitive configurations were then computed for each network, but the TSC weights that were adjusted using FRIWA were removed because their values were manipulated.

Figure 8 shows the average strengths of each adjustable TSC weight over all resistant or all sensitive configurations as an asterisk or diamond, respectively. Each adjustable TSC weight in Figure 8 has 3 asterisk-diamond pairs in 3 different colors (red, blue, and green), illustrating the results from each of the 3 representative networks on a single plot. This figure illustrates closeness in the averages of all the TSC strengths between the resistant and sensitive configurations in all 3 representative networks, demonstrating that no single TSC mediates the therapeutic state by itself. The results of this analysis suggest that all adjustable TSCs contribute to the attainment of the therapeutic state and that no single TSC is determinative of it. This model result suggests that there may not exist a single, individual TSC that can be targeted to alleviate depressive symptoms in all patients (see Discussion).

**Figure 8:**
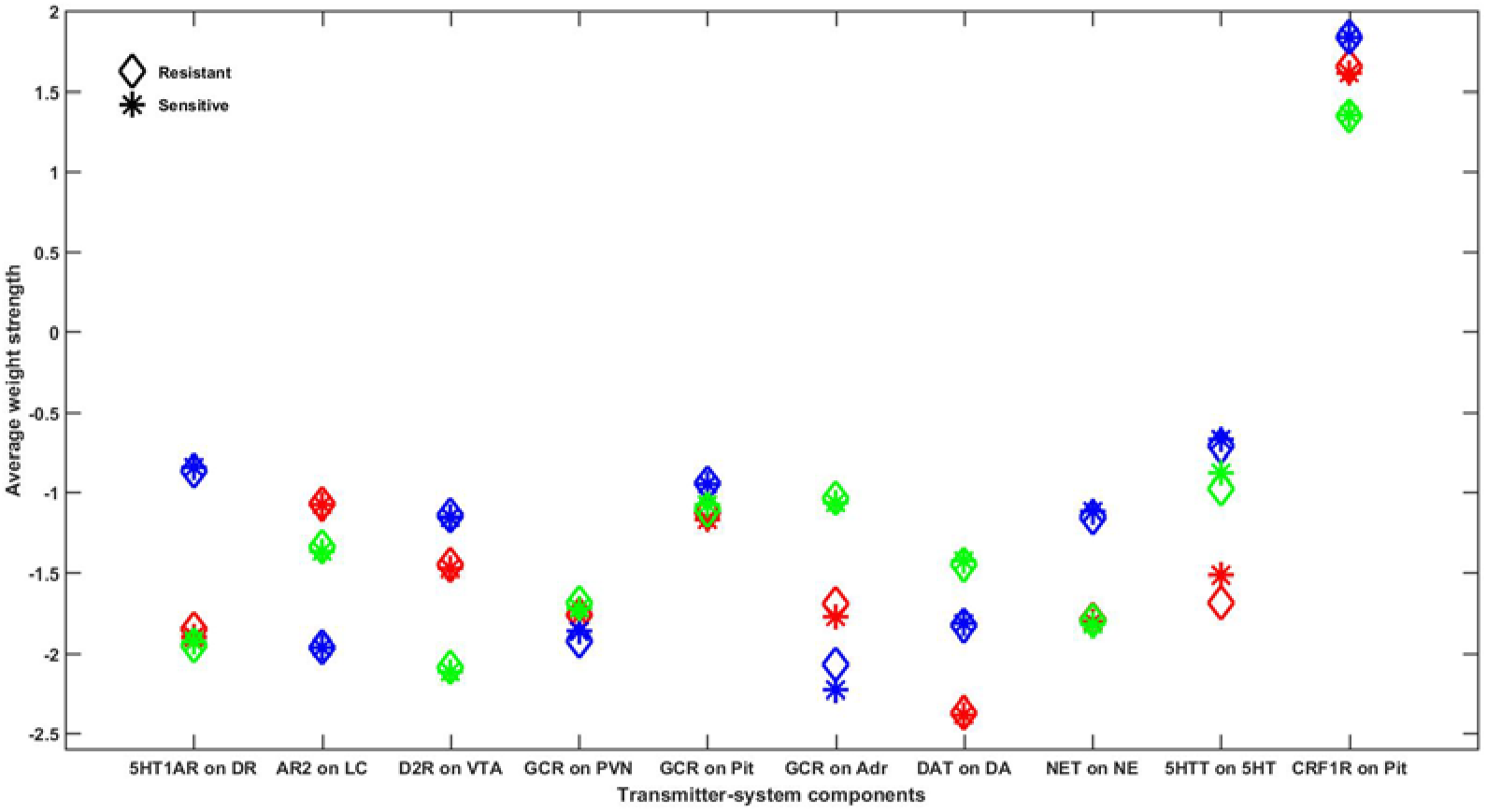
Comparison of average adjustable TSC strengths between resistant and sensitive configurations in all representative networks. Every configuration adapted to chronic SSRI that was also therapeutic (high 5HT and low CORT) at degree 6 was assessed for resistance to adjustments of each of the 10 adjustable TSCs. All TSC strength adjustments that resulted in configurations that were no longer adapted were excluded. Configurations that remained adapted and therapeutic following weight adjustments were determined to be “resistant” and adapted configurations that were no longer therapeutic following TSC adjustments were determined to be “sensitive.” The average strength of each of the 10 adjustable TSCs in all of the configurations, excluding those in which that TSC itself was adjusted, was computed for both the resistant and sensitive configurations and plotted as asterisks or diamonds, respectively. The results for all 3 networks are represented on this single plot using 3 different colors (red, blue, or green) to distinguish between the mean adjustable TSC strengths of each network. Note that the average resistant and sensitive strengths for each adjustable TSC are very close in all 3 networks, illustrating that each individual TSC can provide a contribution to therapeutic resistance, but no single TSC by itself determines the therapeutic state. Abbreviations: 5HT1A receptor, 5HT1AR; α-2 adrenergic receptor, AR2; glucocorticoid receptor, GCR; DA transporter, DAT; NE transporter, NET; 5HT transporter, 5HTT; CRF1 receptor, CRF1R; dorsal raphe, DR; locus coeruleus, LC; ventral tegmental area, VTA; paraventricular nucleus of the hypothalamus, PVN; pituitary gland, Pit; adrenal gland, Adr.

### 3.5 Pairwise Correlations between Adaptable TSC Strengths

The FRIWA analysis considered 1 adjustable TSC at a time. It does not rule out the possibility that therapeuticity may be a property of correlations between pairs of TSCs rather than of single TSCs by themselves. To evaluate this possibility, pairwise correlations between all TSC weights over either all resistant or all sensitive configurations were computed for each representative network, but again, the TSC weights that were adjusted using FRIWA were removed because their values were manipulated. Even at the permissive significance level of p = 0.05, the analysis found no significant pairwise correlations between adjustable TSCs that were consistent over all 3 representative networks in either the resistant or the sensitive configurations.

### 3.6 Temporal-logic Model-checking

As described in previous subsections, the FRIWA and correlation analyses examined a large space of configurations adapted to an SSRI and determined that no single TSC, nor pairs of TSCs, mediates therapeutic resistance by themselves. Neuroadaptation to chronic antidepressant, where each TSC change is incremental, can also be examined as a process to determine if specific states or degrees of neuroadaptation must be reached prior to arriving at therapeutic configurations. Linear temporal-logic (LTL) analysis can be used to elaborate all pathways of sequential TSC-strength adjustments in order to evaluate possible temporal relationships in neuroadaptation (see Methods). This mode of analysis can be used to determine whether certain numbers of TSC-strength adjustments, once attained, will always lead to an adapted and therapeutic state.

The antecedent of the LTL propositions we analyzed specified that a specific TSC had been adjusted 3 times, either up or down (see Methods for details). The antecedent of 3 adjustments was chosen because it was halfway between 0 and 6, the total number of adjustments to which we were limited for technical reasons. The consequent queried whether all subsequent sequences of up to 3 adjustments in any subset of the 10 TSCs would all lead to an adapted and therapeutic state (5HT >= 0.70 and CORT <= 0.70). Each LTL model-check was carried out using each of the 3 networks for 6 total adjustments in all TSCs, and only those model-checking results that were consistent over all 3 networks are reported. All propositions returned false.

We next examined the more easily satisfiable propositions that once a specific TSC had been adjusted 3 times, either up or down, then all subsequent sequences of up to 3 adjustments in any subset of the 10 TSCs would all lead to an adapted and therapeutic state, or the TSC no longer maintains its degree of adjustment. This proposition allowed for the possibility that failure to maintain an adapted and therapeutic state occurred because the specific TSC had not maintained the specified level of adjustment. All of these more satisfiable propositions, however, again returned false, as shown in Table 2.

**Table 2:** Linear temporal-logic (LTL) analysis on the relationship between TSC-adjustments and a therapeutic state. Each LTL analysis was used to evaluate whether 3 steps of sensitization (adjustment up) or desensitization (adjustment down) of a specific TSC during neuroadaptation to chronic SSRI leads, over an ensuing 3 additional adjustments over any subset of the 10 TSCs, either to adapted and therapeutic configurations, or to failure to maintain sensitization or desensitization of the specific TSC. Only the results of LTL model-checks that were in agreement in all 3 networks are reported here. All propositions returned false, meaning that desensitization or sensitization of single adjustable-TSCs for 3 adjustment steps is not determinative of subsequent adapted and therapeutic states.

| Experiment | Condition (at 3 adjustments and beyond) | Leads to 5HT high and CORT low? |
| --- | --- | --- |
| 1 | DR 5HT1AR desensitization | False |
| 2 | LC AR2 desensitization | False |
| 3 | VTA D2R desensitization | False |
| 4 | 5HT 5HTT desensitization | False |
| 5 | NE NET desensitization | False |
| 6 | DA DAT desensitization | False |
| 7 | PVN GCR desensitization | False |
| 8 | Pituitary GCR desensitization | False |
| 9 | Adrenal GCR desensitization | False |
| 10 | Pituitary CRF1R desensitization | False |
| 11 | DR 5HT1AR sensitization | False |
| 12 | LC AR2 sensitization | False |
| 13 | VTA D2R sensitization | False |
| 14 | 5HT 5HTT sensitization | False |
| 15 | NE NET sensitization | False |
| 16 | DA DAT sensitization | False |
| 17 | PVN GCR sensitization | False |
| 18 | Pituitary GCR sensitization | False |
| 19 | Adrenal GCR sensitization | False |
| 20 | Pituitary CRF1R sensitization | False |

The LTL analysis shows that 3 adjustments of any single adjustable TSC weight up or down will not guarantee that a therapeutic state will be reached. All propositions returned false to degree 6, and it is unlikely that they would be true were the LTL analysis extended to greater degrees of adjustment because that would entail additional opportunities for TSC de-adjustment and overall de-adaptation. We did not attempt an exhaustive LTL analysis because the number of potentially relevant propositions that could be tested is simply too many. The result we generated, that 3 adjustments either up (sensitization) or down (desensitization) in no single TSC by itself can lead to states that are all therapeutic within 3 additional adjustments over the other 9 TSCs or itself, supports the FRIWA analysis finding that adjustment of no single weight individually can determine the therapeutic state when the other TSCs can also adjust. These findings impose potential challenges to effective antidepressant drug design (see Discussion).

## 4. Discussion

A consensus is forming around the general idea that multifactorial diseases should be treated with multidrug therapies (Anastasio, 2017; Perry et al., 2015; Xu et al., 2015). It has become common practice to treat psychiatric disorders with combinations of drugs, but the approach has been *ad hoc* and based mainly on clinical trial-and-error (Barowsky and Schwartz, 2006; Zhou et al., 2015). SSRI non-responders are often treated with combinations of two drugs (Trivedi et al., 2006), but systematic evaluation of the relative benefits of different combinations of two or more drugs in depression treatment has not occurred.

The identification of novel, multidrug treatments for depression could proceed either through rational design or brute-force screening of drug combinations. The main challenge associated with rational design is the overwhelming complexity of the neurobiology of depression, which makes it extremely difficult to know *a priori* how any specific drug combination will work. The main challenge associated with brute-force screening is the sheer number of possible drug combinations, which grows geometrically with the number of drugs.

Our model addresses both challenges. It addresses rational design by computationally representing many aspects of the structure and function of the monoaminergic transmitter and stress hormone systems, whose interactions are central in depression neurobiology. It addresses combinatorial explosion by providing the means computationally to screen a large set of drug combinations, and to identify the most promising combinations for clinical evaluation. Addressing these dual challenges is crucial, because treatment with SSRI alone, which is currently the first-line treatment for depression, is less than 40% effective, and efficacy is not greatly increased when SSRI is combined with another drug (Cipriani et al., 2016; Turner et al., 2008; Zhou et al., 2015).

Our previous, initial model of the neurobiology of depression offered an explanation for the low SSRI efficacy rate. That model (the <u>M</u>onoamine or M-model) represented the monoaminergic transmitter systems (and some related transmitter systems) but did not represent the stress hormone system. It did, however, contain 11 adjustable TSCs. It also admitted of many different TSC-strength configurations, and many of those were adaptive in that adaptation error in the presence of simulated, chronic SSRI was lower than initial error. In the M-model, however, only 29% of the configurations adapted to chronic SSRI also elevated 5HT to therapeutic levels (Camacho and Anastasio, 2017). The conclusion was that the clinically observed efficacy of chronic SSRI is low (<40%) because the real monoaminergic neurotransmitter system likewise has many ways to adapt to chronic SSRI, but only some of those ways also elevate 5HT to therapeutic levels.

Our current model (the <u>M</u>onoamine-<u>S</u>tress or MS-model) represents a significant advance over our previous, initial model (the M-model) in several respects. The MS-model represents the interactions of the 3 monoaminergic systems not only with each other but also with the stress hormone system. By adopting a recurrent nonlinear network formalism (see Supplemental Material S3: Details on Model Training), the MS-model represents many more relevant neurobiological details and conforms model behavior to a much larger set of empirical observations.

In the M-model, we analyzed the TSC-strength configurations of a single representative network, but since individuals can vary as well as their TSC-strength configurations, it is necessary to analyze more than 1 representative network. In the MS-model we therefore analyze the TSC-strength configurations of 3 representative networks. In the M-model, we analyzed all TSC-strength configurations reachable in 3 adjustment steps. In the MS-model we double the number of allowed adjustment steps to 6. This represents an increase in the number of configurations analyzed from 11,155 for the M-model (with 11 TSCs but only 1 representative instance) to 382,747 MS-model configurations (with 10 TSCs and 3 representative instances). In the current work we also extend our analytical repertoire to include graphical (distribution/histogram), sensitivity (FRIWA), correlation, and LTL analysis.

The MS-model recapitulates and extends the key idea established in the M-model, that the brain has many ways to adapt to chronic drug administration but not all of those ways will achieve therapeutic goals. An innovation of the current work is the use of histograms to show how the monoamine and CORT levels are distributed over a large set of adapted TSC-strength configurations (Figures 4, 5, and 6). To the extent that the model reflects actual neurobiology, these histograms illustrate how we might expect a chronic drug or drug combination to affect a clinical population of people who differ in their pre-drug, starting configurations and also differ in their subsequent adaptation pathways. Specifically, they illustrate that we should not expect everyone in a depressed population being treated with chronic antidepressants to raise their monoamines, nor to lower their CORT, to therapeutic levels. Computing the average monoamine level over a whole distribution of computationally determined, adapted TSC-strength configurations (Figure 7), is a plausible way to predict the overall, clinical efficacy of chronically administered antidepressant drugs or drug combinations.

The fact that only a subset of the many possible TSC-strength configurations is both adaptive and therapeutic raises the possibility that one, or only a few, of the adjustable TSCs are determinative of the therapeutic state. We used FRIWA analysis (our customized variant of sensitivity analysis) to explore this possibility for single TSCs in all 3 representative networks. We found that some configurations were “resistant” in that they remained adapted and therapeutic despite individual-weight adjustment (IWA) applied to a TSC. Other configurations were “sensitive” in that they did not remain adapted and therapeutic despite IWA applied to a TSC. The average values of individual TSC strengths, however, were essentially the same in resistant versus sensitive configurations (Figure 8) in all 3 networks, indicating that no TSC by itself is determinative of the therapeutic state.

We then used correlation analysis to explore the possibility that pairs of TSCs were determinative of the therapeutic state, and considered all pairs that were significantly correlated at the relatively permissive significance level of p = 0.05. We found that no pair of TSCs was significantly correlated in either the resistant or sensitive configurations over all 3 representative networks, showing that no pairs of TSCs are together determinative of the therapeutic state.

We then used LTL analysis to explore the possibility that a specific TSC, once it reached a certain level of sensitization or desensitization, could guarantee a therapeutic outcome. For technical reasons (see Methods), the number of steps of adjustment we allowed in the strength of each TSC was limited to 6, and our LTL analysis took this constraint into account. For each TSC, we tested the truth or falsehood of two LTL propositions essentially asserting that if a given TSC has reached a certain level of sensitization or desensitization, then the network is guaranteed to arrive at an adapted and therapeutic configuration no matter what the other TSCs do in the remaining adjustment steps. All of these propositions returned false in our LTL analysis (Table 2). What all of these analyses show is that no single TSC, nor pair of TSCs, nor TSCs that have attained a certain degree of adjustment, can determine the therapeutic state. They demonstrate, more generally, that the properties of interest in the model (the level of adaptation, the levels of the 3 monoaminergic transmitters, and the level of CORT) are overdetermined by the 10 TSCs.

In its overdeterminedness (or degeneracy), the MS-model makes contact with other models depicting phenomena at both network and neuronal levels of neurobiological organization. In the vertebrate brain, sensory signals, motor commands, and information in general is represented not by single neurons but by networks of neurons. These representations are overdetermined because the pieces of information are few relative to the number of neurons in the network that represents them. In consequence, the same information can be distributed in many different ways over a network, and neurons in the same network can vary greatly in their response properties in what has been termed a non-uniform distributed representation (Anastasio, 2010, 1991).

Overdetermination of physiological properties by the values of multiple, relevant parameters has also been demonstrated for single neurons and computational models of thereof (Bucher, 2005; Edelman and Gally, 2001; Golowasch et al., 2002; reviewed in Marder and Taylor, 2011). The most well-known case in point was established by Eve Marder and colleagues in their model of the lateral pyloric neuron (LPN) in the lobster somatogastric ganglion (Taylor et al., 2009). In the invertebrate brain, information is sometimes represented not by neural networks but by single, very large (i.e. “giant”) neurons that are identifiable from one animal to another. Experiments revealed that many different ion channels determine the electrophysiological properties of LPNs, and that the parameters that determine the function of ion channels of specific types can vary between LPNs from different decapods (Golowasch et al., 2002; Golowasch and Marder, 1992; MacLean et al., 2005; Schulz et al., 2007, 2006).

The Marder group developed a biophysically detailed model of the LPN and identified 17 parameters (i.e. the conductances and some related properties of 10 ion channel types) that plausibly could vary between individual LPNs. They generated ~6 x 10^5^ different configurations of these 17 parameters and found, after evaluating them all computationally, that 1304 of those configurations endowed the LPN model with the same, realistic set of electrophysiological properties (number of spikes, spike frequency, spike duration, burst duration, etc.). Rather than grouping into specific sets of values, however, the Marder team found that LPN models having equally realistic electrophysiological properties could vary greatly in the values of their 17 defining, ion-channel parameters. By examining parameter value distributions and correlations, they found further that no single parameter nor pair of parameters was determinative of LPN model properties.

Analogously, we find that many different configurations of TSC-strengths can endow the MS-model with the property of adaptation to simulated, chronic SSRI, but we take this a step further and show that the adapted configurations can also differ in other properties, namely in their levels of the monoamines. Overall, these computational studies suggest that there is a large amount of degeneracy in neurobiological systems in general. The MS-model specifically implicates overdeterminedness as a consequence of the complexity inherent in the antidepressant response that poses a challenge to the identification of an antidepressant drug or hormone combination that is effective in all patients (Edelman and Gally, 2001). An approach that recognizes different subtypes of depression, and matches those with monoamine profiles predicted from computational models of the adapted antidepressant response, could provide an effective means of identifying promising drug combinations.

The MS-model identifies drug and hormone combinations that could potentially be more therapeutic than single drugs for patients suffering from specific subtypes of depression. Because the therapeutic effects of chronic antidepressant use have been associated with elevations in monoamine levels and reductions in CORT levels, we were specifically interested in configurations that reproduced these experimentally and clinically observed changes (Lenze et al., 2011; Nikisch et al., 2005). Our model demonstrates that combining an SSRI with certain antipsychotics, atypical antidepressants, or hormones can increase the proportion of adapted states that are also associated with elevations in the monoamines and reductions in CORT.

The MS-model identifies Asenapine, Olanzapine, Bupropion, and Oxytocin as adjuncts to SSRI therapy that hold particular promise. Precedent for the use of these adjuncts has been established clinically. Physicians in practice frequently resort to the augmentation of SSRIs with antipsychotics such as Asenapine and Olanzapine in SSRI-monotherapy non-responders (Boulton et al., 2010; Han et al., 2013). The atypical antidepressant Bupropion has also been found to relieve depressive symptoms in SSRI non-responders and is widely used clinically, either by itself or in conjunction with SSRI (Leuchter et al., 2008; Trivedi et al., 2006; Zisook et al., 2006). Intranasal administration of the hormone Oxytocin combined with chronic SSRI also can be more effective in treating depressed patients than SSRI by itself (Scantamburlo et al., 2015).

Results from the MS-model are consistent with these general findings in that pairing SSRI with either Asenapine, Olanzapine, Bupropion, or Oxytocin increases the number of adapted states with elevated monoamines and reduced CORT over that observed with SSRI alone (Figures 5 and 6). Furthermore, the triples composed of SSRI/Asenapine/Oxytocin or of SSRI/Bupropion/Olanzapine further increase the proportion of adapted states associated with therapeutic changes in monoaminergic neurotransmitter and CORT levels in the MS-model (Figures 5 and 6). The model suggests that these triples could be clinically useful.

To predict the efficacy rates of potentially useful drug combinations, the average levels of each of the monoamines, over all configurations of all 3 representative networks, that were adapted to SSRI combined with 1 or 2 other selected agents were computed (Figure 7). By illustrating the average level of each monoamine in response to chronic administration of SSRI in combination with selected drugs or hormones in the MS-model, the heatmap in Figure 7 can help guide physicians in their choices of drugs and hormones with which to augment SSRI therapy. Combinations that included an SSRI and an antipsychotic tended to be located at the upper range of monoamine levels (closer to the therapeutic reference vector [0.70 0.70 0.70]), and combinations that included Bupropion tended to elevate average NE and DA levels closer to the therapeutic reference vector. The average monoamine levels of combinations that included Oxytocin hormone were also located near the therapeutic reference vector.

This analysis can potentially fine-tune combined pharmacotherapy for depression, as different subtypes of depression have previously been shown to respond best to elevations in the levels of the different monoamines (Malhi et al., 2005; Parker, 2000). Specifically, patients with non-melancholic, melancholic, or psychotic depression have been observed to respond best to antidepressants that elevate 5HT, NE, or DA, respectively (Guelfi et al., 1995; Malhi et al., 2005, 2002; Parker et al., 1992). Drug combinations that included Trazodone (an SNRI), Olanzapine (an antipsychotic), and GBR-12935 (a DAT antagonist) tended to raise 5HT, NE, and DA levels, respectively, in the model.

The computational screen predicts potentially adverse as well as beneficial combinations. Notably, combinations that include both an SSRI and an MAOI were found to increase monoaminergic neurotransmitter levels well beyond that of other combinations. This MS-model result is in line with the clinical finding that combining these two drugs can result in toxic elevations of the monoamines (Boyer and Shannon, 2005; Remick and Froese, 1990; Shrier et al., 2000).

## Conclusion

Our overall MS-modeling strategy is both rational and brute force. It is rational in that the model incorporates, both in its structure and its function, many aspects of the known interactions between the monoamine and stress-hormone systems—two systems that are central to depression neurobiology. It is brute force in that it can be used to screen large numbers of drug combinations computationally, and to predict the distributions of monoamine levels that could be expected in a diverse patient population in which each individual can adapt to the chronic drug regimen in their own unique way. The MS-model can be used to explore neuroadaptive configurations and to propose drug/hormone combinations that could be therapeutic for a higher proportion of patients than single-drugs by themselves. It could also be used to direct antidepressant drug-design toward specific depressive subtypes, by predicting the average levels of each of the three monoamines that would be expected in a patient population following adaptation to chronic administration of various drug combinations.

## Competing Interests

The authors declare that the research was conducted in the absence of any commercial or financial relationships that could be construed as a potential conflict of interest.

## Funding

The authors declare no competing financial interests.

## Acknowledgments

We thank Ignacia Caviedes, Haven Comeaux, Kate Hamblen, Emily Hamm, Neena Joshi, Mahak Lalani, Diana Masolak, Cassandra Mora, and Katherine Zitello for their help in compiling the database of experimental facts on which the MS-model is based. We thank Allegra Domel for her contribution to the computational pruning analysis. We also thank Shivali Patel for her computational contribution to the construction of the full model-diagram.

## Supplementary Material

Supplementary information is available at the publisher’s website.

