## Supplementary Material for "Computational Analysis of Therapeutic Neuroadaptation to Chronic Antidepressant in a Model of the Monoaminergic Neurotransmitter and Stress Hormone Systems"

### Unpublished Preprint: Privileged Communication

#### Supplemental Material:

This document serves as a supplement to the manuscript, **Computational Analysis of Therapeutic Neuroadaptation to Chronic Antidepressant in a Model of the Monoaminergic Neurotransmitter and Stress Hormone Systems**.

#### Contents

***Text***

**S1**: Details on Structure Connections 1

**S2**: Truth-table Justification 5

**S3**: Details on Model Training 20

**S4**: Pruning Methods 20

**S5**: Details on Temporal-logic Model-checking Procedure 21

***Figures***

**Supplemental Figure 1**: Complete model structure diagram 26

**Supplemental Figure 2**: Agreement between desired (i.e., target) and actual outputs is high either after training but before pruning, or after pruning and re-training 27

***Tables***

**Supplemental Table 1:** Complete model truth-table 28

**Supplemental Table 2:** Canonical model weights 28

**Supplemental Table 3:** Adjustable TSCs____________________________________________30

***References***_______________________________________________________________32

**S1: Details on Structure Connections**

The structure matrix includes the subset of interactions between units in the model that are known from experimental observation. Inhibitory connections were given a −1, excitatory connections were given a +1, and connections of unknown polarity were given a 2 in the structure matrix. If the polarity was unknown by the literature, it was set by training algorithm.

**Inputs**

**SSRIs** inhibit the 5HT transporter protein, so there is an inhibitory connection from SSRI to the 5HT transporter (Sanchez, Bergqvist et al. 2003, Nemeroff and Owens 2004). **Nomifensine** blocks the NE and DA transporters, and weakly blocks the 5HT transporter, so there are inhibitory connections from Nomifensine to NET, DAT, and 5HTT (Schacht and Heptner 1974, Brogden, Heel et al. 1979, Tatsumi, Groshan et al. 1997). **Reboxetine** potently blocks the NE transporter, so there is an inhibitory connection from Reboxetine to NET (Hajos, Fleishaker et al. 2004). **Trazodone** inhibits the 5HT transporter, is an agonist at the 5HT1A receptor, an antagonist of 5HT2A, 5HT2C, A1, and A2 receptors (Waldmeier 1982, Cusack, Nelson et al. 1994, Krege, Goepel et al. 2000). Asenapine is an agonist at 5HT1A receptors and an antagonist at 5HT2A, 5HT2C, A1, A2, D1, D2, D3, and D4 receptors (Balsara, Jadhav et al. 2005, Odagaki, Toyoshima et al. 2005, Corporation 2009, Shahid, Walker et al. 2009, Stahl 2009, Stahl 2009). **Aripiprazole** is an agonist at 5HT1A, 5HT2C, D2, D3, and D4 receptors, an antagonist at 5HT2A receptors, and interacts with A1, A2, B1, and B2 receptors with unknown polarity (Keck and McElroy 2003, Kroeze, Hufeisen et al. 2003, Shapiro, Renock et al. 2003). **Bupropion** inhibits the NE and DA transporters, so it has inhibitory projections to NET and DAT (Waldmeier 1982, Cooper, Wang et al. 1994, Stahl, Pradko et al. 2004). Humans metabolize **Quetiapine** to N-Desalkyl quetiapine (Nquet) so we simulated the effects of Quetiapine in the model by giving Quetiapine all the targets of both Quetiapine and Nquet (DeVane and Nemeroff 2001, Jensen, Rodriguiz et al. 2008). Quetiapine inhibits the NE transporter and 5HT2A, 5HT2C, A1, A2, D1, D2 and D3 receptors and is an agonist of 5HT1A receptors (DeVane and Nemeroff 2001, Jensen, Rodriguiz et al. 2008). **Pramipexole** (PPX) is an agonist at D2, D3 and D4 receptors (Mierau, Schneider et al. 1995). GBR is a DA transporter antagonist (TOCRIS , Andersen 1989). **Clozapine** is an agonist at 5HT1A receptors, is an antagonist at 5HT2A, 5HT2C, A1, D2, and D4 receptors, and has unknown polarity at A2, D1, and D3 receptors (Roth , Meltzer 1994, Newman-Tancredi, Chaput et al. 1996). **Ketamine** is an antagonist at NMDA receptors (Vollenweider and Kometer 2010). **Reserpine** depletes monoamines by inhibiting the activity of the vesicular monoamine transporter 2 (VMAT2), which transports monoamines to the cell membrane for release into the extracellular space (Scherman and Henry 1984, Rudnick, Steiner-Mordoch et al. 1990). Because VMAT2 is not an element in our model, Reserpine sends inhibitory projections directly to 5HT, NE and DA. **Venlafaxine** is a selective serotonin-norepinephrine reuptake inhibitor (SNRI) with some antagonistic effects at the DA transporter

(Roseboom and Kalin 2000, Wellington and Perry 2001). **Desipramine** is a tricyclic antidepressant that strongly inhibits the NE transporter and inhibits the 5HT and DA transporters with weaker affinity (Berti and Shore 1967). It is also an antagonist of the 5HT2A (Wander, Nelson et al. 1986). **CP96345** (CP) is a neurokinin-1 receptor antagonist (Fong, Yu et al. 1992). **Gepirone** is an agonist of 5HT1A receptors (Blier and Ward 2003). **RU28362** (RU) is a glucocorticoid receptor agonist (Sernia and Thomas 1994). **Org34850** (Org) is a glucocorticoid receptor antagonist (Reynolds, Saunders et al. 2015). **Oxytocin** is a hormone input that projects to oxytocin receptors in the model (Gimpl and Fahrenholz 2001). **Dexamethasone** is a corticosteroid that acts as an agonist at glucocorticoid receptors but not mineralocorticoid receptors (Bamberger, Bamberger et al. 1995, Reul, Gesing et al. 2000, Pariante and Miller 2001). **WAY100635** (WAY) is a selective 5HT1A receptor antagonist (Fletcher, Forster et al. 1996). Monoamine Oxidase Inhibitors (**MAOI**) block the activity of monoamine oxidase, which is an enzyme that mediates the breakdown of the monoamines (Stein 1960, Remick and Froese 1990). M617 is a galanin receptor 1 (galR1) agonist (Blackshear, Yamamoto et al. 2007). The CRF1R antagonist input has an inhibitory projection to the CRF1 receptor. **Haloperidol** is a dopamine D2 receptor antagonist (Schotte, Janssen et al. 1993). **Olanzapine** is dopamine D2 receptor antagonist and 5HT2A receptor antagonist (Pilowsky, Busatto et al. 1996). **Clonidine** is an alpha-2 receptor agonist (Unnerstall, Kopajtic et al. 1984). **Yohimbine** is an alpha-2 receptor antagonist (Perry and U'Prichard 1981). **DR lesion**, **LC lesion**, **VTA lesion**, **PVN lesion**, **Amygdala lesion**, **Hippocampus lesion**, **PFC lesion** and Adrenalecotmy are inhibitory to DR, LC, VTA, PVN, Amygdala, Hippocampus, PFC, the adrenal gland, respectively. The **stress** response begins with activation of the HPA axis, so the stress input sends an excitatory projection to the PVN (Sapolsky 2000, Gold and Chrousos 2002, de Kloet, Joels et al. 2005). **Exogenous ACTH** is excitatory to ACTH receptors (Kitay, Holub et al. 1959). **Exogenous CRF** is excitatory to CRF1 and CRF2 receptors (Merchenthaler 1984).

**<u>Bias</u>**

Each unit has a bias of unknown polarity.

**<u>Regions</u>**

Each region has an excitatory (+1) connection to the units it secretes. The **DR** secretes 5HT, glutamate, galanin, and NK1 (Melander, Hokfelt et al. 1986, Johnson 1994, Santarelli, Gobbi et al. 2001, Guiard, Guilloux et al. 2007, Soiza-Reilly and Commons 2011, Gagnon and Parent 2014, Liu, Zhou et al. 2014). The **LC** secretes NE, galanin, and NK1 (Holets, Hokfelt et al. 1988, Jordan, Kermadi et al. 1995, Pieribone, Xu et al. 1995, Maubach, Martin et al. 2002, Kawa, Barde et al. 2016). The **VTA** secretes DA and glutamate (Stuber, Hnasko et al. 2010, Root, Mejias-Aponte et al. 2014). The **amygdala** secretes glutamate, CRF, AVP, and NK1 (Buijs and Swaab 1979, Merchenthaler 1984, Curtis, Bello et al. 2002, Roberto, Schweitzer et al. 2004, Singewald, Chicchi et al. 2008). The **PFC** secretes glutamate (Bagley and Moghaddam 1997, Geisler, Derst et al. 2007, Albert, Vahid-Ansari et al. 2014). The **PVN** secretes CRF, AVP, oxytocin, and NK1 (Antoni, Fink et al. 1990, Yamashita, Kasai et al. 1991, Makara 1992, Emiliano, Cruz et al. 2007, Rodaros, Caruana et al. 2007, Bulbul, Babygirija et al. 2011, Feetham and Barrett-Jolley 2014). The **hippocampus** secretes glutamate (Bagley and Moghaddam 1997). The **pituitary gland** secretes ACTH and the **adrenal gland** produces cortisol (Kitay, Holub et al. 1959, Butler, Clarke et al. 1969, Seiden and Brodish 1971, Loose, Do et al. 1980).

**Enzymes and Substrates**

Each enzyme or substrate has either an excitatory (+1) or inhibitory (−1) connection with its corresponding enzyme or substrate. The 5HT transporter is inhibitory on 5HT, the NE transporter is inhibitory on NE and DA, and the DA transporter is inhibitory on DA (Iversen 1971, Carboni, Tanda et al. 1990, Giros, Wang et al. 1994). In order to represent the 5HT synthesis pathway, there is an excitatory connection from tryptophan to tryptophan hydroxylase (TPH), an excitatory connection from TPH to 5-hydroxytryptamine (5HTP), 5HTP to 5HT-decarboxylase (5HTDC), from 5HTDC to 5HT, from 5HT to N-acetylserotonin O-methyltransferase (ASMT), and from ASMT to melatonin (Schott, Nicolai et al. 2010). Synthesis of NE and DA were represented with an excitatory connection from tyrosine to tyrosine hydroxylase (TH), an excitatory connection from TH to levadopa (L-DOPA), an excitatory connection from L-DOPA to aromatic L-amino acid decarboxylase (L-AADC), an excitatory connection from L-AADC to DA, an excitatory connection from DA to dopamine beta-hydroxylase (DBH), and an excitatory connection from DBH to NE (Cosentino, Marino et al. 2015). There is an inhibitory connection from monoamine oxidase (MAO) to 5HT, NE and DA to represent how MAO is responsible for the breakdown of the monoamines (Edmondson, Mattevi et al. 2004).

**Transmitters**

The neurotransmitters and hormones all have excitatory connections (+1) to their corresponding receptors to the model. 5HT projects to 5HT1A, 5HT1B, 5HT2A, and 5HT2C receptors. NE projects to A1, A2, B1, and B2 receptors. DA projects to D1, D2, D3, and D4 receptors. Glutamate projects to alpha-amino-3-hydroxy-5-methyl-4-isoxazolepropionic acid (AMPA) and N-methyl-D-aspartate (NMDA) receptors. Galanin projects to galR1 and galR2. AVP projects to vasopressin receptor 1a (V1AR). CRF projects to CRF1 and CRF2 receptors. Cortisol projects to glucocorticoid (GC) and mineralocorticoid (MC) receptors. Oxytocin projects to oxytocin receptors. NK1 projects to NK1 receptors. ACTH projects to melanocortin 2 (MC2) receptors. GABA projects to GABA receptors.

**Receptors**

Receptors have either excitatory (+1), inhibitory (−1), or unknown polarities on the brain regions and neurotransmitters. To account for the effect of presynaptic receptors, known presynaptic receptors had projections to neurotransmitters whose levels are regulated by the presynaptic receptor.

5HT1A receptors are inhibitory on DR, LC, PFC, and hippocampus neurons (Azmitia, Gannon et al. 1996, Blier, Pineyro et al. 1998, Celada, Puig et al. 2004, Santana, Bortolozzi et al. 2004, Puig, Artigas et al. 2005, Lanzenberger, Baldinger et al. 2013). 5HT1A receptors are also present on PVN neurons, but their overall polarity on PVN neurons has not been determined conclusively due to their presence on GABAergic interneurons impinging on PVN neurons (Pan and Gilbert 1992, Dinan 1996). Presynaptic 5HT1A receptors have been found to inhibit GABA release (Koyama, Matsumoto et al. 2002). 5HT2A receptors have been found to be inhibitory on LC but excitatory on the amygdala, PFC, and hippocampus (Haddjeri, de Montigny et al. 1997, Xu and Pandey 2000, Szabo and Blier 2002, Santana, Bortolozzi et al. 2004, Bombardi 2011). A1 receptors are excitatory on DR, amygdala, and PVN neurons (Baraban and Aghajanian 1980, Clement, Gemsa et al. 1992, Itoi, Suda et al. 1994, Cecchi, Khoshbouei et al. 2002). A2 receptors are inhibitory autoreceptors on LC neurons, and also reside with unknown polarities on VTA, amygdala, PFC, hippocampus, and PVN neurons (Cedarbaum and Aghajanian 1977, Talley, Rosin et al. 1996). AR2 receptors have also been found presynaptically to decrease NE, galanin, and NK1 levels (Cedarbaum and Aghajanian 1977, Kuraishi, Hirota et al. 1985, Tsuda, Yokoo et al. 1989, Bertolino, Vicini et al. 1997). B1 and B2 receptors are excitatory on LC, PFC and hippocampus neurons (Rainbow, Parsons et al. 1984). D1 receptors exist in the PFC with unknown polarity (Pirot, Godbout et al. 1992, Hall, Sedvall et al. 1994, Chen and Yang 2002). D2 receptors are inhibitory autoreceptors on VTA neurons, are excitatory on amygdala, DR, and PFC neurons, and exist in the PVN with unknown polarity (Ferre and Artigas 1993, Hall, Sedvall et al. 1994, Haj-Dahmane 2001, Rosenkranz and Grace 2002, Brady and O'Donnell 2004, Succu, Sanna et al. 2007). D3 receptors are present in the PFC and PVN with unknown polarity (Levesque, Diaz et al. 1992, Succu, Sanna et al. 2007). D4 receptors are present in the PFC with unknown polarity (Primus, Thurkauf et al. 1997). AMPA and NMDA receptors have been found in the DR, LC, VTA, amygdala, PFC, and PVN, and vary in polarity depending on whether they are present on GABAergic interneurons or not, so their polarities were not fixed (Petralia and Wenthold 1992, Martin, Blackstone et al. 1993, Huntley, Vickers et al. 1994, de Kock, Cornelisse et al. 2006). galR1s are inhibitory galanin receptors, and they have been found to inhibit DR, LC, VTA, amygdala, and PVN neurons (Perez, Basile et al. 2002, Hawes and Picciotto 2004, Hawes, Brunzell et al. 2005). galR2s are excitatory on DR and hippocampus neurons (Hawes and Picciotto 2004, Elliott-Hunt, Pope et al. 2007). V1ARs have been found on amygdala and hippocampus neurons, and are excitatory on DR neurons and the pituitary gland (Ostrowski, Lolait et al. 1992, Rood and Beck 2014). CRF1 receptors are inhibitory on DR neurons, but are excitatory on LC, VTA, amygdala, PVN, and the pituitary gland (Van Pett, Viau et al. 2000, Kirby, Freeman-Daniels et al. 2008, Reyes, Valentino et al. 2008, Wanat, Hopf et al. 2008, Ji, Fu et al. 2013, Sparta, Hopf et al. 2013, Baiamonte, Valenza et al. 2014, Reyes, Bangasser et al. 2014). CRF2 receptors are excitatory on DR and VTA neurons (Wang, You et al. 2007, Spannuth, Hale et al. 2011, Wood, Zhang et al. 2013). GC receptors have been found on DR, LC, VTA, and hippocampus neurons with unknown polarity (Makara and Haller 2001, Makino, Smith et al. 2002, Lanfumey, Mongeau et al. 2008, Vincent and Jacobson 2014). They are inhibitory to PVN neurons, the pituitary gland, adrenal gland, and they decrease expression of the 5HT transporter (Fumagalli, Jones et al. 1996, Pariante and Lightman 2008, Heydendael and Jacobson 2010). They have been found to increase activity and/or expression of TPH, MAO, and TH (Clark, Pai et al. 2005, Lindley, She et al. 2005, Ou, Chen et al. 2006, Heydendael and Jacobson 2009). Mineralocorticoid receptors have been found in the hippocampus with unknown polarity (Herman, Watson et al. 1993, Patel, Lopez et al. 2000, Yau,

Noble et al. 2001). They also mediate negative feedback on the HPA axis through inhibitory effects on the PVN, pituitary gland, and adrenal gland (Ratka, Sutanto et al. 1989, Patel, Lopez et al. 2000, Ladd, Huot et al. 2004). Oxytocin receptors have been found in the DR, LC, VTA, hippocampus, and pituitary gland with unknown polarity (Elands, Beetsma et al. 1988, Breton, Pechoux et al. 1995, Gimpl and Fahrenholz 2001, Yoshida, Takayanagi et al. 2009). They are inhibitory in the amygdala (Buijs and Swaab 1979, Huber, Veinante et al. 2005). NK1 receptors are inhibitory on DR neurons, and have been found on LC, VTA, amygdala, PFC, and PVN neurons with unknown polarity (Tooney, Au et al. 2000, Conley, Cumberbatch et al. 2002, Rigby, O'Donnell et al. 2005, Renoldi and Invernizzi 2006, Gobbi, Cassano et al. 2007, Guiard, Guilloux et al. 2007, Haddjeri and Blier 2008, Feetham and Barrett-Jolley 2014). MC2 receptors are excitatory on the adrenal gland (Papadimitriou and Priftis 2009). GABA receptors have been found to be inhibitory on DR neurons, and have also been found on LC, VTA, amygdala, PFC, hippocampus, and PVN neurons, as well as the pituitary gland with unassigned polarities due to the presence of interneurons modulating their effects on the corresponding neurons (Baraban and Aghajanian 1980, Chang, Tran et al. 1980, Biggio, Corda et al. 1981, Roland and Sawchenko 1993, Van Bockstaele and Pickel 1995, Hatfield, Spanis et al. 1999, Cai, Flores-Hernandez et al. 2002, Katsurabayashi, Kubota et al. 2003, Puig, Santana et al. 2004, Tan, Zhong et al. 2004, Cornelisse, Van der Harst et al. 2007, Kreft and Zorec 2008, Bulbul, Babygirija et al. 2011, Jin, Bhandage et al. 2014, Kudo, Konno et al. 2014).

**S2: Truth-table Justification**

The MS model was trained on baseline values and to reproduce data on the effects of neuron activations, transmitter, enzyme, and hormone levels of acute receptor blockade, lesions, and other experimental manipulations. All of the manipulations (drug or hormone administration, chemical lesioning, etc.) will be referred to as “inputs.” For the purposes of training the model the training data were assembled into an input-output table or “truth table” in which the inputs are acute drug administrations, and the outputs are changes from baseline firing-rate of the monoaminergic neurons expressed as a percentage. Data on the effects of inputs on model unit activations were obtained through an extensive literature search that compiled findings from multiple groups using a broad range of experimental methods. Training data were for 40 single inputs and 25 combination inputs. Each row of the truth table represents the results of one or more actual experiments where output levels were measured in response to one or more inputs. Inputs are either present or absent (0 or 1) and outputs are assigned integer values between 0.30 (lowest) to 0.70 (highest). Output findings from the literature and their corresponding integer values are shown in the table below. All outputs bound between 0 and 1 by the sigmoidal squashing function.

| 0.30 | 0.40 | 0.50 | 0.60 | 0.70 |
|---|---|---|---|---|
| Decrease maximally | Decrease moderately | No change | Increase moderately | Increase maximally |

Determinations of “moderate” versus “maximal” changes were made based on both the qualitative and quantitative data that was available. When more than one finding was available on a particular input-output relationship, a consensus was reached using the available data. If neurotransmitter level changes differed in different brain regions, the change in the level of the neurotransmitter in the prefrontal cortex (PFC) was used in the truth table. If whole brain neurotransmitter level change was available, then the effect of the input on whole brain neurotransmitter level was used in the truth table. If neurotransmitter level change data for the whole brain or PFC was not available, then the change in the neurotransmitter level of the brain regions that were available was used to determine the truth-table value with that input. This subjective approach was necessary to account for differences between research labs in experimental methods, drug dosages and routes of administration, and levels of quantification. The majority of the findings were derived from rodent studies but some were obtained from horse, monkey, cat, cow, fish, squirrel and human studies.

**SSRI**

The first single-input is an **SSRI**, which consolidates the findings associated with acute administration of several different SSRIs. Although different SSRIs have different low-affinity off-target effects, all of them have strong affinity for the 5HT transporter, which is the only structure connection from the SSRI input included in the model (Tatsumi, Groshan et al. 1997, Sanchez, Bergqvist et al. 2003). Binding of the SSRI to the 5HT transporter leads to a doubling of extracellular 5HT, so the output for 5HT with acute SSRI was set to 0.60 (Invernizzi, Belli et al. 1992, Koch, Perry et al. 2002, Calcagno, Guzzetti et al. 2009). Because acute administration of SSRIs other than fluoxetine does not significantly change NE and DA levels, the truth table values for these neurotransmitters were set to 0.50 (Bymaster, Zhang et al. 2002, Koch, Perry et al. 2002).

The increase in extracellular 5HT associated with acute SSRI administration has been associated with increased binding to the 5HT1A autoreceptor on DR neurons, decreasing the firing activity of these neurons (Calcagno, Guzzetti et al. 2009). The Blier group and others have found that the DR neuron firing rate decreases by about 65% with acute SSRI, so the truth table value for this output was set to 0.40 (de Montigny, Chaput et al. 1990, Czachura and Rasmussen 2000, Chernoloz, El Mansari et al. 2012). The Blier group found that acute SSRI administration decreases the firing rate of LC neurons by 45% and decreases the firing rate of VTA neurons by 41%, so the truth table values for the LC and VTA firing rates with acute SSRI were also set to 0.40 (Chernoloz, El Mansari et al. 2009, Dremencov, El Mansari et al. 2009, Chernoloz, El Mansari et al. 2012). fMRI studies indicate that amygdala activity decreases moderately, while cortical activity does not change with acute SSRI administration (Mayberg, Brannan et al. 2000, Kennedy, Evans et al. 2001, Takahashi, Yahata et al. 2005, Murphy, Norbury et al. 2009). The Murphy lab found that amygdala activity decreases moderately in response to neutral faces with acute SSRI, and the Mayberg lab found that PFC activity increases after 6-weeks of SSRI treatment but not after acute (1-week)

SSRI treatment. The truth table values for the amygdala and PFC with acute SSRI are set to 0.40 and 0.50, respectively.

Acute SSRI administration has been found to stimulate the rodent HPA axis by increasing CRF, ACTH, and cortisol levels while also increasing PVN activity (Jensen, Jessop et al. 1999, Wieczorek, Schulz et al. 2001, Hesketh, Jessop et al. 2005). Specifically, 30 minutes of subcutaneous cannula citalopram administration maximally increases cortisol and ACTH levels (Jensen, Jessop et al. 1999). This lab also found that PVN activity increases moderately as measured by the percentage of c-Fos immunoreactive cells in the PVN. The same lab did not find a significant difference in CRF mRNA with acute citalopram treatment; however, another lab found that acute citalopram moderately increases CRF levels (Moncek, Duncko et al. 2003). The Moncek lab also found that acute citalopram produces very large increases in cortisol and ACTH levels. The truth table values for ACTH and cortisol with acute SSRI was both set to 0.70. The truth table value for CRF with acute SSRI was set to 0.60 to reflect the moderate increase in CRF found by the Moncek lab and the finding that cortisol and ACTH levels increase maximally. Oxytocin levels have been found to moderately increase with acute SSRI, while arginine-vasopressin (AVP) levels have been found to stay the same (Hesketh, Jessop et al. 2005). We set the oxytocin target to 0.60 and the AVP target to 0.50 with acute SSRI. Acute fluoxetine injection moderately increases galanin mRNA levels, so the truth-table value for galanin with acute SSRI was set to 0.60 (Kuteeva, Wardi et al. 2008).

**Nomifensine**

The second single-input is **Nomifensine**, a reuptake blocker of the NE and DA transporter proteins, with slight affinity for the 5HT transporter (Samanin, Bernasconi et al. 1975, Brogden, Heel et al. 1979, Tatsumi, Groshan et al. 1997). The Blier group found that acute administration of Nomifensine increases DR neuron activity by 50%, decreases VTA neuron activity by 39%, and decreases LC neuron activity by 71% (Katz, Guiard et al. 2010). The truth-table values for the acute effect of Nomifensine on DR, LC and VTA neurons were therefore set to 0.60, 0.40, and 0.40, respectively. The Masana lab found that acute Nomifensine maximally increases DA levels, but the Samanin lab found that acute Nomifensine did not significantly change DA levels (Samanin, Bernasconi et al. 1975). The Carboni lab found that acute Nomifensine increases DA levels by a very large amount (Carboni, Imperato et al. 1989). The Butcher lab found a moderate increase in DA levels with acute Nomifensine (Butcher, Fairbrother et al. 1988). Because some labs report a maximal increase in DA with acute Nomifensine while others do not detect any change in DA levels, the truth-table value for DA with acute Nomifensine was set to 0.60 to reflect a moderate increase in DA.

**Reboxetine**

**Reboxetine** is a selective NE transporter blocker (Hajos, Fleishaker et al. 2004). The Blier group found that acute Reboxetine administration decreases LC firing rate by 68% without affecting DR firing rate (Szabo and Blier 2001). They also found that acute Reboxetine administration decreases VTA neuron firing rate by 31% (Katz, Guiard et al. 2010). The truth-table values for acute Reboxetine for DR, LC, and VTA were therefore set to 0.50, 0.40, and 0.40, respectively. Acute Reboxetine administration doubles NE levels and produces a moderate increase in DA levels while producing no change in 5HT levels, corresponding to values of 0.60, 0.60, and 0.50 in the truth table for these neurotransmitters, respectively (Page and Lucki 2002). One fMRI study in humans shows that acute Reboxetine administration moderately decreases amygdala response to neutral stimuli, so the truth-table value for the amygdala with acute Reboxetine was set to 0.40 (Onur, Walter et al. 2009). Acute Reboxetine administration in humans has been found to moderately increase ACTH levels in two studies using male volunteers, so the truth-table value for ACTH with acute Reboxetine was set to 0.60. Acute Reboxetine administration has also been shown to moderately increase cortisol levels in two different studies using male human volunteers, so the truth-table value for cortisol with acute Reboxetine was set to 0.60 (Hennig, Lange et al. 2000, Schule, Baghai et al. 2004).

**Trazodone**

**Trazodone** is a multifunctional drug that blocks the 5HT transporter and interacts with multiple monoaminergic receptors (Stahl 2009, Stahl 2009). The Blier lab found that acute Trazodone administration decreases DR neuron firing by 65%, increases LC neuron firing by 25%, and does not alter VTA neuron firing (Ghanbari, El Mansari et al. 2010, Ghanbari, El Mansari et al. 2012). We set the target output values for DR, LC, and VTA with acute Trazodone to 0.40, 0.60, and 0.50, respectively. Acute Trazodone administration has been found to double 5HT levels without changing NE levels, so the truth-table values for these were set to 0.60 and 0.50, respectively (Rowbotham, Jones et al. 1984, Pazzagli, Giovannini et al. 1999).

**Asenapine**

**Asenapine** is an antipsychotic drug that interacts with multiple monoaminergic receptors (Franberg, Wiker et al. 2008, Ghanbari, El Mansari et al. 2009). The Blier group found that acute Asenapine administration decreases DR neuron firing by about 30% without affecting the firing rates of the LC or VTA (Oosterhof, El Mansari et al. 2015). We set the truth table values for DR, LC, and VTA to 0.40, 0.50, and 0.50. Acute Asenapine administration in rats has been found to moderately increase 5HT, NE, and DA levels (Franberg, Marcus et al. 2009). The truth table outputs for 5HT, NE, and DA with acute Asenapine were all set to 0.60.

**Aripiprazole**

**Aripiprazole** is an antipsychotic drug with strong affinity for DA and 5HT receptors (Shapiro, Renock et al. 2003). The Blier group found that acute Aripiprazole administration increases DR neuron firing rate by 48%, without affecting the firing rates of LC or VTA neurons (Chernoloz, El Mansari et al. 2009). The truth-table values for the DR, LC and VTA were set to 0.60, 0.50 and 0.50, respectively. Acute Aripiprazole administration has been found to moderately increase DA levels in experiments done by Zocchi et al and Li et al without affecting 5HT, NE, or cortisol levels (Li, Ichikawa et al. 2004, Zocchi, Fabbri et al. 2005, Assie, Carilla-Durand et al. 2008). The truth-table values for 5HT, NE, DA, and cortisol were set to 0.50, 0.50, 0.60, and 0.50 with acute Aripiprazole, respectively.

**Bupropion**

**Bupropion** is an antidepressant drug that blocks the NE and DA transporter proteins (Cooper, Wang et al. 1994, Stahl, Pradko et al. 2004). The Blier group found that acute Bupropion administration doubles DR neuron firing rate, decreases LC neuron firing rate by half, and does not change VTA neuron firing rate (El Mansari, Ghanbari et al. 2008). Another group found that acute Bupropion moderately decreases LC and VTA neuron firing rate (Cooper, Wang et al. 1994). The truth-table values for the DR, LC and VTA were set to 0.60, 0.40 and 0.40, respectively, with acute Bupropion administration. Acute Bupropion administration has been found to moderately increase DA and NE levels without affecting 5HT levels (Piacentini, Clinckers et al. 2003). The truth-table values for 5HT, NE, DA were set to 0.50, 0.60, and 0.60 with acute Bupropion.

**Quetiapine**

**Quetiapine** is an antipsychotic drug with multiple receptor and transporter targets, including the dopamine D2 receptor, the 5HT2A receptor, and the alpha-1 adrenergic receptor (AR1) (DeVane and Nemeroff 2001, Jensen, Rodriguiz et al. 2008). The Blier group found that acute Quetiapine administration decreases the DR neuron firing rate by 43% and increases the LC neuron firing rate by 40% (Chernoloz, El Mansari et al. 2012). The truth-table values for the DR and LC with acute Quetiapine were set to 0.40 and 0.60, respectively. Another group found that acute Quetiapine moderately increases VTA firing rate, so the truth-table value for the VTA with acute Quetiapine was set to 0.60 (Werkman, Olijslagers et al. 2004). Denys et al found that acute Quetiapine moderately increases 5HT and DA levels in the PFC, while Silverstone et al found that acute Quetiapine has no effect on 5HT levels, but moderately increases NE and DA levels (Denys, Klompmakers et al. 2004, Silverstone, Lalies et al. 2012). The truth table values for 5HT, NE, and DA with acute Quetiapine were all set to 0.60.

**Pramipexole**

**Pramipexole** (PPX) is a D2, D3, and D4 receptor agonist (Mierau, Schneider et al. 1995). The Blier group found that acute PPX administration does not change DR neuron firing rate, decreases LC neuron firing rate by 33%, and decreases VTA neuron firing rate by 40% (Chernoloz, El Mansari et al. 2009). The truth-table values for the DR, LC and VTA were therefore 0.50, 0.40 and 0.40 with acute PPX administration.

**GBR12909**

**GBR12909** (GBR) is a DA transporter blocker (TOCRIS , Andersen 1989, Singh 2000). The Blier group found that acute GBR administration does not change DR or LC neuron firing rate but decreases VTA neuron firing rate by 26% (Katz, Guiard et al. 2010). Another group found a similar decrease of about 25% in VTA neuron firing rate with acute GBR (Choong and Shen 2004). The truth-table values for the DR, LC and VTA were set to 0.50, 0.50 and 0.40, respectively, with acute

GBR administration. Three groups have found moderate increases in DA levels with acute GBR, so the truth-table value of DA with acute GBR was set to 0.60 (Rothman, Mele et al. 1991, Choong and Shen 2004, Masana, Bortolozzi et al. 2011).

**<u>Clozapine</u>**

**Clozapine** is an antipsychotic drug that targets many serotonergic, noradrenergic, and dopaminergic receptors (Meltzer 1994). Acute administration of Clozapine has been found to moderately decrease the firing activity of DR neurons, moderately increase the firing rate of LC neurons, moderately increase the firing rate of VTA neurons, and moderately increase the firing activity of PFC neurons (Souto, Monti et al. 1979, Sprouse, Reynolds et al. 1999, Chen and Yang 2002, Gao 2007). Another group found that acute Clozapine completely inhibits DR firing activity, however, the dose that was used was much higher than the standard rodent dose (Gallager and Aghajanian 1976). The truth-table values for DR, LC, VTA, and PFC were set to 0.40, 0.60, 0.60, and 0.60, respectively. Acute Clozapine administration has been found by Zocche et al to maximally increase NE levels and moderately increase DA levels. Masana et al also found that Clozapine moderately increases DA levels. Lee et al 2001 found that Clozapine moderately increases cortisol levels in awake humans, so the truth table value for NE was set to 0.70 and the truth table values for DA and cortisol were set to 0.60 (Zocchi, Fabbri et al. 2005, Masana, Bortolozzi et al. 2011). One group has found that Clozapine decreases 5HT release in the nucleus accumbens, while another found that Clozapine increases 5HT in the nucleus accumbens and the PFC (Ferre and Artigas 1995, Ichikawa, Kuroki et al. 1998). Because these two groups found opposing effects of Clozapine on 5HT, and a third group found that 5HT does not change in the PFC with acute Clozapine, we used a truth-table value of 0.50 for 5HT with acute Clozapine (Zocchi, Fabbri et al. 2005).

**<u>Ketamine</u>**

**Ketamine** is an antagonist at N-methyl-D-asparatate (NMDA) receptors (Hall and Murdoch 1990). The Blier group found that acute Ketamine administration does not change DR or VTA neuron firing rate but increases LC neuron firing rate by 23% (El Iskandrani, Oosterhof et al. 2015). The truth-table values for the DR, LC and VTA were set to 0.50, 0.60, and 0.50, respectively, with acute Ketamine administration. Razoux et al found that Ketamine moderately increases PFC neuron activity and moderately increases extracellular glutamate. Stone et al also found that Ketamine moderately increases anterior cingulate glutamate, so the truth-table value for the PFC and glutamate were set to 0.60 (Razoux, Garcia et al. 2007, Stone, Dietrich et al. 2012, Pehrson and Sanchez 2014, Bjorkholm, Franberg et al. 2015). Nishitani et al found that Ketamine moderately increases PFC 5HT levels, so the truth table value for 5HT was set to 0.60 with acute Ketamine. Stone et al and Lindeforsa et al both found that Ketamine has no effect on GABA levels, so the truth table value for GABA was set to 0.50 with acute Ketamine. Glisson et al found that acute Ketamine had no effect on rabbit whole brain NE or NE levels in any of the brain areas examined, so the truth table value for NE with acute Ketamine was set to 0.50. Glisson et al also found that DA levels did not change in rabbit whole brain with Ketamine, but did find a moderate increase in thalamus and hypothalamus DA. Lindeforsa et al found that acute Ketamine moderately increases DA levels in rat PFC. Because whole brain neurotransmitter level change is the standard criterion for the truth table, the truth-table value for DA with Ketamine was set to 0.50.

**Reserpine and Reserpine/Bupropion**

**Reserpine** depletes monoamines by inhibiting the activity of the vesicular monoamine transporter 2 (VMAT2), which transports monoamines to the cell membrane for release into the extracellular space (Scherman and Henry 1984, Rudnick, Steiner-Mordoch et al. 1990). Because VMAT2 is not an element in our model, Reserpine sends inhibitory projections directly to 5HT, NE and DA. Reserpine maximally decreases 5HT, NE, and DA levels, so the truth-table values for 5HT, NE, and DA were all set to 3 (Cooper, Wang et al. 1994). Cooper et al found that Reserpine did not affect LC neuron firing rate, so the truth table value for LC with acute Reserpine was set to 0.50. Baraban et al found that Reserpine moderately increases DR activity, then suppresses DR activity after about 30 minutes (Baraban, Wang et al. 1978, Baraban and Aghajanian 1980). Because we are interested in the immediate, acute effect of Reserpine, we set the truth-table value for DR with acute Reserpine to 0.60. An injection of Reserpine in cows produces no change in cortisol levels, so the truth-table value for cortisol was set to 0.50 for Reserpine (Bauman, Collier et al. 1977). The combination of Reserpine and Buproprion produces no change in the firing rate of LC neurons, so the truth-table value for LC with Reserpine/Bupropion was set to 0.50 (Cooper, Wang et al. 1994).

**Venlafaxine**

**Venlafaxine** is a selective serotonin-norepinephrine reuptake inhibitor (SNRI) (Roseboom and Kalin 2000). The Blier group found that Venlafaxine decreases DR firing rate by 47% and decreases LC firing rate by 21%, so the truth-table values for DR and LC were both set to 0.40 (Gartside, Umbers et al. 1997). Because acute Venlafaxine administration moderately increases the firing activity of the PFC in a c-fos study, we set the truth-table value for PFC with Venlafaxine to 0.60 (Higashino, Ago et al. 2014). Higashino et al found that acute Venlafaxine administration moderately increases 5HT in the PFC, moderately increases NE in the PFC, and maximally increases DA in the PFC and Beyer et al found that acute Venlafaxine has no effect on 5HT levels but maximally increases NE levels in the PFC (Beyer, Boikess et al. 2002, Higashino, Ago et al. 2014). The truth-table value for 5HT was set to 0.60 to reflect a moderate increase due to Venlafaxine's 5HT transporter inhibition. The truth-table values for NE and DA were both set to 0.70 to reflect the large increases in the levels of these neurotransmitters with acute Venlafaxine observed by the Higashino and Beyer groups.

**Desipramine**

**Desipramine** is a tricyclic antidepressant with strong affinity for the NE transporter and weaker affinity for the 5HT and DA transporters as well as affinity for various monoaminergic receptors (Berti and Shore 1967). DR firing was found to moderately decrease with acute Desipramine, so the truth-table value for DR was set to 0.40 with acute Desipramine (Gartside, Umbers et al.

1997). Acute Desipramine also moderately decreases LC neuron activity, so the truth-table value for LC was set to 0.40 (Scuvee-Moreau and Dresse 1979). Acute administration of Desipramine was found by Beyer et al to maximally increase NE levels without changing 5HT levels, and Kreiss et al also found that Desipramine has no effect on 5HT levels (Kreiss and Lucki 1995, Beyer, Boikess et al. 2002). Higashino et al found no change in 5HT levels with acute Desipramine, but found a maximal increase in NE levels and a moderate increase in DA levels (Kreiss and Lucki 1995, Higashino, Ago et al. 2014). The truth-table value for 5HT was set to 0.50, for NE was set to 0.70, and for DA was set to 0.60. Acute Desipramine was found to have no effect on PFC activity, so this truth-table value was set to 0.50 (Higashino, Ago et al. 2014). Desipramine injection in humans moderately increases blood cortisol levels after 30 minutes, so the truth-table value for cortisol with Desipramine was set to 0.60 (Asnis, Halbreich et al. 1985).

**CP-96345 and CP-96345/Stress**

**CP-96345** (CP) is a neurokinin-1 receptor antagonist (Fong, Yu et al. 1992). The Blier group found that acute administration of CP increases the firing rate of DR neurons by 46% but does not affect the firing rate of LC neurons, so the truth-table values for DR and LC were set to 0.60 and 0.50 for the DR and LC with acute CP, respectively (Conley, Cumberbatch et al. 2002, Haddjeri and Blier 2008). Lejeune et al found that NK1 receptor antagonists moderately enhance VTA firing rate, so the truth-table value for VTA with acute CP was set to 0.60. Lejeune et al also found that NK1 receptor antagonists have no effect on 5HT levels, so the 5HT value with acute CP was set to 0.50. Renoldi et al found that acute CP has no effect on NE levels, but moderately increases DA levels in the PFC of gerbils. Lejeune et al also found a moderate increase in DA levels with acute NK1 receptor antagonists. The truth table value for NE was set to 0.50 and for DA was set to 0.60 with acute CP (Lejeune, Gobert et al. 2002, Renoldi and Invernizzi 2006, Guiard, Guilloux et al. 2007). Renoldi et al also found that the combination of NK1 antagonists and stress leads to no change in NE or DA levels (Renoldi and Invernizzi 2006). The truth-table values for NE and DA with CP/Stress were set to 0.50.

**Gepirone, Gepirone/Stress, and Gepirone/Dexamethasone**

**Gepirone** is an agonist of 5HT1A receptors (Blier and Ward 2003). The Blier group and others have found that acute administration of 5HT1A agonists maximally decrease DR neuron firing rate, so the truth-table value of DR with Gepirone was set to 0.30 (VanderMaelen, Matheson et al. 1986, Blier and de Montigny 1987, Blier and de Montigny 1990). 5HT1A receptor agonists have been found to maximally decrease 5HT levels by multiple groups, so the truth table value for 5HT with acute Gepirone was set to 0.30 (Rutter, Gundlah et al. 1994, Dawson, Nguyen et al. 2002). 5HT1A receptor agonists have also been found to moderately increase CRF, ACTH, and cortisol release in rodents, so the truth-table values for CRF, ACTH, and cortisol with acute Gepirone were set to 0.60 (Pan and Gilbert 1992, Matheson, Knowles et al. 1997). The combination of Gepirone and stress also moderately increases cortisol levels, so the cortisol level in the truth-table with acute Gepirone/stress was set to 0.60 (Matheson, Knowles et al. 1997). The combination of Gepirone/Dexamethasone moderately decreases cortisol levels, so the cortisol level in the truth-table with acute Gepirone/Dexamethasone was set to 0.40 (Matheson, Knowles et al. 1997).

## RU-28362

**RU-28362** (RU) is a glucocorticoid receptor agonist (Sernia and Thomas 1994). Administration of RU alone does not change ACTH levels, so the truth-table value for ACTH with acute RU was set to 0.50 (Hinz and Hirschelmann 2000).

#### Org-34850 and Org-34850/Stress

**Org-34850** (Org) is a glucocorticoid receptor antagonist (Reynolds, Saunders et al. 2015). Glucocorticoid receptors mediating negative feedback on the HPA axis are present in the hypothalamus, pituitary gland, and adrenal gland (among others) (Loose, Do et al. 1980, Morimoto, Morita et al. 1996, Ozawa, Ito et al. 1999). Acute Org administration by itself has been found by Spiga et al 2007 and Spiga et al 2008 to have no effect on cortisol levels, so the truth-table value for cortisol with acute Org was set to 0.50. Acute **Org/Stress** was also found by this group to have no effect on cortisol levels, so the truth-table value for cortisol with acute Org/Stress was set to 0.50.

#### Oxytocin and Oxytocin/Stress

**Oxytocin** is an input that projects to oxytocin receptors in the model (Gimpl and Fahrenholz 2001). Oxytocin administration to rats has been shown to double DA levels, so the truth-table value for DA was set to 0.60 (Melis, Melis et al. 2007). Oxytocin by itself as well as **Oxytocin/Stress** had no effect on PVN CRF mRNA, so the CRF value with Oxytocin and with Oxytocin/Stress was set to 0.50 (Bulbul, Babygirija et al. 2011). Oxytocin has been found to moderately decrease amygdala response to fearful stimuli (faces and scenery), so the truth-table value for amygdala with Oxytocin/Stress was set to 0.40 (Kirsch, Esslinger et al. 2005). One study in rats from 1984 found that Oxytocin does not have a significant effect on ACTH levels, however, because several other groups since 1984 have found that Oxytocin moderately decreases ACTH in both humans and monkeys, the truth-table value for ACTH with Oxytocin was set to 0.40 (Gibbs, Vale et al. 1984, Chiodera and Coiro 1987, Parker, Buckmaster et al. 2005). It has also been found that Oxytocin moderately decreases cortisol levels, so the truth-table value for cortisol with Oxytocin was set to 0.40 (Legros, Chiodera et al. 1984).

#### Dexamethasone

**Dexamethasone** is a corticosteroid with significant affinity for glucocorticoid receptors but not mineralocorticoid receptors in vivo (Bamberger, Bamberger et al. 1995, Reul, Gesing et al. 2000, Pariante and Miller 2001). Rush et al found that Dexamethasone maximally suppresses cortisol levels, so the truth-table value for cortisol with Dexamethasone was set to 0.30 (Rush, Giles et al. 1996). Kovacs et al found that Dexamethasone maximally suppresses PVN CRF mRNA, so the truth-table value for CRF with Dexamethasone was set to 0.30 (Kovacs and Mezey 1987). Hohnloser et al found that acute Dexamethasone administration moderately decreases ACTH levels, so the truth-table value for ACTH with Dexamethasone was set to 0.40 (Hohnloser, Von Werder et al. 1989). Kovacs et al found that acute Dexamethasone moderately decreases AVP mRNA in the PVN, so the truth-table value for AVP with Dexamethasone was set to 0.40 (Kovacs and Makara 1988). C-Fos mRNA responses in the PVN and pituitary gland were found to be moderately decreased with acute Dexamethasone, so the truth-table values for the PVN and pituitary gland were set to 0.40 (Karssen, Meijer et al. 2005). Dexamethasone has been shown to interact with the monoamines. Specifically, Dexamethasone administration has been shown to moderately increase the levels of 5HT precursors and 5HT, so the truth-table values for Trp, 5HTP, and 5HT were set to 0.60 with Dexamethasone (Maes, Meltzer et al. 1995, Tsubota, Adachi et al. 1999, Clark, Flick et al. 2008). Dexamethasone has also been found to moderately increase the levels of extracellular DA but moderately decrease extracellular NE, so the truth-table values for DA and NE were set to 0.60 and 0.40, respectively (Stene, Panagiotis et al. 1980, Tsubota, Adachi et al. 1999). Oxytocin levels were unaffected by Dexamethasone, so the truth table value for oxytocin with Dexamethasone was set to 0.50 (Fink, Robinson et al. 1988).

**WAY-100635 and WAY-100635/Gepirone**

**WAY-100635** (WAY) is a selective 5HT1A receptor antagonist (Fletcher, Forster et al. 1996). Administration of WAY has been shown to reverse the inhibitory effects of 5HT1A receptor agonists on DR firing rate, so the truth-table value for DR with **WAY/Gepirone** was set to 0.50 (Fletcher, Forster et al. 1996). WAY/Gepirone together has also been shown to prevent the rise in ACTH observed with Gepirone, so the truth-table value for ACTH with WAY/Gepirone was set to 0.50 (Fletcher, Forster et al. 1996). 5HT1AR antagonists by themselves moderately increase extracellular 5HT, so the truth-table value for 5HT with WAY was set to 0.60 (Arborelius, Nomikos et al. 1996).

**MAOI**

Monoamine Oxidase Inhibitors (**MAOI**) block the activity of monoamine oxidase, which is an enzyme that mediates the breakdown of the monoamines (Stein 1960, Remick and Froese 1990). The Blier group found that acute MAOI administration moderately decreases DR and LC neuron firing without affecting VTA neuron firing, so the truth-table values for DR and LC were set to 0.40 and the truth table value for VTA was set to 0.50 with MAOI (Blier and de Montigny 1985, Chenu, El Mansari et al. 2009). Acute MAOI administration has been shown to produce maximal increases in all 3 of the monoamines, so the truth-table values for 5HT, NE and DA were set to 0.70 (Butcher, Fairbrother et al. 1990, Celada and Artigas 1993, Kitaichi, Inoue et al. 2006).

**M-617**

Galanin is a 29 amino acid neuropeptide that is widely distributed in the central nervous system and co-secreted by DR and LC neurons (Tatemoto, Rokaeus et al. 1983). Agonists to the galR1 receptor (**M-617**) have been shown to have a pro-depressive effect, which is believed to be related to the inhibitory effect of galR1 receptors on DR and LC neurons (Sevcik, Finta et al. 1993, Larm, Shen et al. 2003, Wang, Li et al. 2016). Acute M-617 administration has been found to moderately decrease DR and LC firing rate and moderately decrease 5HT and NE levels (Jacobs, Wise et al. 1974, Azmitia and Segal 1978, Seutin, Verbanck et al. 1989, Sevcik, Finta et al. 1993, Yoshitake, Reenila et al. 2003, Hawes, Brunzell et al. 2005, Mazarati, Baldwin et al. 2005). We therefore set the truth-table values for DR, LC, 5HT, and NE to 0.40. It has also been found that M-617 administration produces moderate increases in the activities of the amygdala and PVN as measured by increases in c-fos expression in these regions, so the truth-table values for these regions were set to 0.60 (Blackshear, Yamamoto et al. 2007).

**CRF1R Antagonist and CRF1R antagonist/Stress**

CRF1 receptor antagonists (**CRF1R antagonist**) block the CRF1 receptor to decrease HPA axis activity (Holsboer and Ising 2008). Specifically, it has been found by multiple labs that CRF1R antagonists moderately decrease ACTH and cortisol release, so the truth-table values for ACTH and cortisol with Antalarmin were set to 0.40 (Broadbear, Winger et al. 2004, Jutkiewicz, Wood et al. 2005, Ising and Holsboer 2007). It has also been found that acute administration of **CRF1R antagonist/Stress** moderately increases ACTH and cortisol, so the truth-table values for ACTH and cortisol for Antalarmin/stress was set to 0.60 (compared to +100% with stress alone) (Deak, Nguyen et al. 1999, Jutkiewicz, Wood et al. 2005).

**Haloperidol and Haloperidol/Bupropion**

**Haloperidol** is a dopamine D2 receptor antagonist (Schotte, Janssen et al. 1993). Subcutaneous haloperidol administration in rats produced no change in PFC NE or DA levels, so the truth table values for NE and DA with acute Haloperidol were set to 0.50 (Li, Perry et al. 1998). The combination of Haloperidol and Bupropion produced no change in the firing rate of VTA neurons, so the truth-table value for VTA with Haloperidol/Bupropion was set to 0.50 (Cooper, Wang et al. 1994).

**Olanzapine**

**Olanzapine** is dopamine D2 receptor antagonist and 5HT2A receptor antagonist (Pilowsky, Busatto et al. 1996). Subcutaneous Olanzapine administration in rats moderately increases PFC NE and DA levels, so the truth table values for NE and DA with acute Olanzapine were set to 0.60 (Li, Perry et al. 1998).

**Clonidine**

**Clonidine** is an α-2 receptor agonist (Unnerstall, Kopajtic et al. 1984). Administration of Clonidine has been found to moderately decrease LC neuron firing in rodents, and moderately decrease NE levels in humans (Veith, Best et al. 1984, Jacobs 1986). The truth-table values for LC and NE with acute Clonidine were set to 0.40.

**Yohimbine and Yohimbine/Stress**

**Yohimbine** is an alpha-2 receptor antagonist (Perry and U'Prichard 1981). Yohimbine administration in rats moderately elevates NE levels in the amygdala, so the truth-table value for

NE with acute Yohimbine was set to 0.60 (Khoshbouei, Cecchi et al. 2002). The combination of Yohimbine and Stress maximally increase NE levels in the amygdala, so the truth-table value for NE with acute Yohimbine/Stress was set to 0.70 (Khoshbouei, Cecchi et al. 2002). The combination of Yohimbine/Stress was also found to moderately elevate galanin levels, so the truth-table value for galanin with Yohimbine/Stress was set to 0.60 (Khoshbouei, Cecchi et al. 2002).

<u>**DR lesion/LC lesion/VTA lesion**</u>

The Blier group did a series of experiments where they lesioned one of the monoaminergic nuclei, then observed the change in the firing activity of the other two monoaminergic regions in order to determine how they influence one another. The results of these experiments are included in the truth table. **Lesioning DR** neurons maximally decreases 5HT levels, **lesioning LC** neurons maximally decreases NE levels, and **lesioning VTA** neurons maximally decreases DA levels (Guiard, El Mansari et al. 2008). The truth-table values of each were set to 0.30, respective to each lesion and neurotransmitter. DR lesion moderately increases LC and VTA firing rates (Haddjeri, de Montigny et al. 1997, Guiard, El Mansari et al. 2008, Ito, Shimogawa et al. 2014). The truth-table values for both LC and VTA with DR lesion were set to 0.60. LC lesion moderately increases VTA firing without changing DR firing rate (Gobbi, Cassano et al. 2007, Guiard, El Mansari et al. 2008). The truth-table values for VTA and DR with LC lesion were set to 0.60 and 0.50, respectively. VTA lesion moderately decreases DR firing rate and moderately increases LC firing rate (Guiard, El Mansari et al. 2008, Guiard, El Mansari et al. 2008). The truth table values for DR and LC with VTA lesion were set to 0.40 and 0.60, respectively.

<u>**Stress**</u>

The **stress** response begins with activation of the HPA axis, so the **stress** input sends an excitatory projection to the PVN (Sapolsky 2000, Gold and Chrousos 2002, de Kloet, Joels et al. 2005). Stress leads to maximal elevation of PVN activity as measured by induction of c-fos mRNA expression (Cullinan, Herman et al. 1995). Stress also leads to maximal increases in the activity of the anterior pituitary gland and adrenal gland as measured by maximal blood flow increases after acute stress and c-fos induction (Goldman 1963, Yang, Koistinaho et al. 1989). The truth-table values for PVN, anterior pituitary, and adrenal gland with acute stress were set to 0.70. Stress maximally increases plasma CRF, ACTH, and cortisol, so the truth-table value for CRF, ACTH and cortisol with stress were set to 0.70 (Goldman 1963, Zimmermann and Critchlow 1967, Harbuz and Lightman 1989). NE levels have been found to moderately increase in response to stressful stimuli (Galvez, Mesches et al. 1996, Hatfield, Spanis et al. 1999). LC neuron activity has been found to moderately increase in response to stress (Abercrombie and Jacobs 1987, Buffalari and Grace 2007). Stress has also been shown to moderately increase tryptophan, 5HTP, 5HT and DA levels in many different brain regions (Thierry, Fekete et al. 1968, Abercrombie, Keefe et al. 1989, Kawahara, Yoshida et al. 1993, Summers, Kampshoff et al. 2003). We set the truth-table values for tryptophan, 5HTP, 5HT, NE and DA to 0.60. Acute stress moderately increases DR neuron firing rate in rodents, and VTA firing rate in cats, so the truth-table values for DR and VTA were set to 0.60 (Trulson and Preussler 1984, Bambico, Nguyen et al. 2009). Acute stress moderately increases glutamate levels, so the truth-table value for glutamate was set to 0.60 (Reznikov, Grillo et al. 2007). Acute stress moderately increases AVP and oxytocin levels, so the truth-table values for AVP and oxytocin were set to 0.60 (Hesketh, Jessop et al. 2005). The amygdala and hippocampus are moderately activated in response to stress while the PFC is moderately inhibited, so the truth-table values for the amygdala, hippocampus, and PFC were set to 0.60, 0.60 and 0.40, respectively (Sakanaka, Shibasaki et al. 1986, Van de Kar, Piechowski et al. 1991, Chen, Fenoglio et al. 2006, Alexander, Hillier et al. 2007, Qin, Hermans et al. 2009). Acute stress moderately elevates melatonin levels, so the truth-table value for melatonin was set to 0.60 (Lynch, Eng et al. 1973, Vollrath and Welker 1988). Foot-shock stress causes a moderate decrease in GABA in rats, so this truth-table value was set to 0.40 (Biggio, Corda et al. 1981). Acute immobilization-stress produces no change in galanin levels in the amygdala, so the truth-table value for galanin with stress was set to 0.50 (Khoshbouei, Cecchi et al. 2002).

**Adrenalectomy, Adrenalectomy/Stress and Adrenalectomy/Dexamethasone**

With the **adrenalectomy** (ADX) input, the adrenal gland truth-table value was set to 0.30 (Iacobone, Albiger et al. 2008). ADX has been found to maximally decrease plasma cortisol levels, so the truth-table value for cortisol with ADX was set to 0.30 (Butler, Clarke et al. 1969). This results in a maximal rise in CRF and ACTH as well as a moderate rise in AVP, so the truth table values for CRF and ACTH were set to 0.70 for AVP was set to 0.60 with adrenalectomy (Vernikos-Danellis 1965, Fink, Robinson et al. 1988, Unno, Wu et al. 1998, Iacobone, Albiger et al. 2008). Because subsequent PVN and pituitary-gland firing rates moderately increase with ADX, we set the truth-table values for PVN and pituitary gland to 0.60 (Kitay, Holub et al. 1959, Wynn, Harwood et al. 1985, Kasai and Yamashita 1988). Oxytocin levels were unaffected by adrenalectomy, so the truth table value for oxytocin with adrenalectomy was set to 0.50 (Fink, Robinson et al. 1988). The combination of ADX/Stress has been found to maximally increase ACTH levels in female rats, so the truth-table value for ACTH with ADX/Stress was set to 0.70 (Vernikos-Danellis 1965). The combination of ADX/Dex has been found to moderately decrease ACTH and AVP levels, so the truth-table values of ACTH and AVP with ADX/Dex were set to 0.40 (Fink, Robinson et al. 1988).

**Exogenous ACTH**

**Exogenous ACTH** projects to ACTH receptors (Kitay, Holub et al. 1959). Intramuscular injections of ACTH in horses leads to a moderate increase in cortisol levels after 2-4 hours, so the truth-table value for cortisol with exogenous ACTH was set to 0.60 (Thorn, Forsham et al. 1950, Larsson, Edqvist et al. 1979). ACTH has been shown to increase adrenal gland activity, so the truth-table value for adrenal gland with exogenous ACTH was set to 0.60 (Yang, Koistinaho et al. 1990).

**Exogenous CRF**

**Exogenous CRF** projects to CRF1 and CRF2 receptors in the structure matrix (Merchenthaler 1984). Exogenous CRF administration in humans has been found to moderately increase plasma

ACTH and cortisol levels (Hermus, Pieters et al. 1984). The truth-table values for ACTH and cortisol with exogenous CRF were both set to 0.60.

**PVN, amygdala, hippocampus, and PFC lesion**

**Lesions** to the **PVN**, **amygdala**, **hippocampus** and **PFC** all result in maximal decreases in PVN, amygdala, hippocampus, and PFC, respectively (Chang, Tran et al. 1980). The truth-table values for PVN, amygdala, hippocampus, and PFC, with PVN lesion, amygdala lesion, hippocampus lesion, and PFC lesion, respectively, were all set to 0.30. PVN lesion maximally reduces CRF levels and moderately reduces oxytocin levels, so the truth-table values for CRF and oxytocin with PVN lesion were set to 0.30 and 0.40, respectively (Bruhn, Plotsky et al. 1984, Antoni, Fink et al. 1990). Basal ACTH levels have also been found to moderately decrease with a PVN lesion, so the truth-table value for ACTH was set to 0.40. PVN lesion has been found to maximally decrease melatonin levels, so the truth-table value for melatonin with PVN lesion was set to 0.30 (Klein, Smoot et al. 1983). Adrenal gland activity has been found to moderately decrease with PVN lesion, so the truth-table value for adrenal gland was set to 0.40 (Makara, Stark et al. 1981).

**PVN, amygdala, hippocampus, and PFC lesion and Stress**

The combination of **PVN lesion and stress** results in a moderate increase in CRF, ACTH, and cortisol (Makara, Stark et al. 1981, Bruhn, Plotsky et al. 1984, Makara 1992). The truth table values for these were therefore set to 0.60. **Amygdala lesion and stress** has been shown to lead to a moderate increase in CRF, ACTH, and cortisol, so the truth table values for these were also set to 0.60 (Sakanaka, Shibasaki et al. 1986, Van de Kar, Piechowski et al. 1991, Feldman, Conforti et al. 1994). However, **lesions of the PFC and hippocampus combined with stress** lead to maximal elevations of ACTH and cortisol (Jacobs, Wise et al. 1974, Herman, Schafer et al. 1989, Jacobson and Sapolsky 1991, Diorio, Viau et al. 1993, Herman, Cullinan et al. 1995). We set the truth-table values for ACTH and cortisol with PFC lesion/Stress and Hippocampus lesion/stress to 0.70.

**PVN lesion/Gepirone**

The combination of PVN lesion and a 5HT1A receptor agonist prevented the rise in ACTH observed with the agonist by itself, so the truth-table value for ACTH with **PVN lesion/Gepirone** was set to 0.50 (Bluet Pajot, Mounier et al. 1995).

**Dexamethasone/CRF**

The **Dexamethasone/CRF** test is widely used in depression research to detect HPA axis dysfunction (Rush, Giles et al. 1996, Kunugi, Ida et al. 2006, Heim, Mletzko et al. 2008). CRF is injected after pretreatment with Dexamethasone (Kunugi, Ida et al. 2006). In normal control subjects, ACTH and cortisol secretion does not increase, but in stressed subjects, this test shows moderate increases in ACTH and cortisol secretion (Hohnloser, Von Werder et al. 1989, Kunugi, Ida et al. 2006, Heim, Mletzko et al. 2008). We therefore set the truth-table values for ACTH and cortisol with Dexamethasone/CRF to 0.50, and the truth-table values for ACTH and cortisol with

**Dexamethasone/CRF/stress** to 0.60.

**SSRI/Stress**

Acute restraint stress with acute SSRI has been found to moderately increase AVP levels, so the truth-table value for AVP with SSRI/stress was set to 0.60 (Hesketh, Jessop et al. 2005). Oxytocin and CRF levels moderately increase with SSRI/stress, so the truth-table values for oxytocin and CRF with SSRI/Stress were set to 0.60 (Hesketh, Jessop et al. 2005). The combination of SSRI/Stress has been found to produce a maximal increase in ACTH and cortisol, so the truth-table values for ACTH and cortisol were set to 0.70 with SSRI/Stress (Hesketh, Jessop et al. 2005). SSRI/Stress has been found to maximally increase NE levels, so the truth-table value for NE with SSRI/stress was set to 0.70 (Page and Abercrombie 1997).

**SSRI/WAY-100635**

The combination of an SSRI and a 5HT1AR antagonist (**SSRI/WAY**) has been found to maximally increase 5HT levels, so the truth-table value for SSRI/WAY was set to 0.70 (Arborelius, Nomikos et al. 1996). The combination of an SSRI and WAY has been found to produce no change in DR firing rate from baseline, so the truth-table value for DR with SSRI/WAY was set to 0.50 (Arborelius, Nomikos et al. 1995, Hajos, Gartside et al. 1995).

**SSRI/Bupropion**

Co-administration of an SSRI and Bupropion (**SSRI/Bupropion**) has been found by the Blier group to double DR firing rate and decrease LC firing rate by 60% (Ghanbari, El Mansari et al. 2010). The truth-table values for DR and LC with SSRI/Bupropion were set to 0.60 and 0.40, respectively. SSRI/Bupropion have been found to moderately increase 5HT, NE and DA levels, so the truth-table values for 5HT, NE and DA with acute SSRI/Bupropion were set to 0.60 (Li, Perry et al. 2002).

**SSRI/Aripiprazole**

The Blier group found that the combination of an SSRI and Aripiprazole (**SSRI/Aripiprazole**) produces no significant change in the firing of DR or VTA neurons, but decreases LC neuron firing by 26%. (Chernoloz, El Mansari et al. 2009). The truth table values for DR, LC, and VTA with SSRI/Aripiprazole were set to 0.50, 0.40, and 0.50, respectively.

**SSRI/Quetiapine**

The Blier group found that the combination of an SSRI and Quetiapine (**SSRI/Quetiapine**) produces a 65% decrease in DR neuron firing and a 27% increase in LC neuron firing (Chernoloz, El Mansari et al. 2012). The truth table values for DR and LC with SSRI/Quetiapine were therefore set to 0.40 and 0.60, respectively.

**Reboxetine/Stress**

The combination of Reboxetine and fearful stimuli (fearful faces) has been found to moderately increase amygdala activity in fMRI experiments (Onur, Walter et al. 2009). The truth-table value for amygdala with Reboxetine/Stress was set to 0.60. The combination of Reboxetine and stress in rodents has been found to produce no change in 5HT levels, maximally increases NE levels, and moderately increase DA levels (Page and Lucki 2002). The truth-table values for 5HT, NE and DA with Reboxetine/Stress were set to 0.50, 0.70, and 0.60, respectively.

**S3: Details on Model Training**

The model was trained on 66 different input patterns. One of these was the “no drug” baseline levels of all the outputs. Our baseline output for 0 net input was set at 0.50 for all output units. This is because neurotransmitter and hormone levels are proportional to the firing rates of the neurons that produce them (Chergui, Suaud-Chagny et al. 1994).

Thirty-nine of the input patterns were single inputs (drug, hormone, or lesion). Twenty-four input patterns were combinations of 2 inputs (e.g., Escitalopram (an SSRI) and Aripiprazole (an antipsychotic drug) and 1 input pattern was a combination of 3 inputs (Dexamethasone (a steroid drug), CRF, and stress). Desired output-unit responses were obtained from the literature in the form of assigned integer values between 0.30 (lowest) to 0.70 (highest) and are described in the previous section and in the main text.

**S4: Pruning Methods**

We were interested in pruning the networks in order to eliminate unneeded non-structure connections, and to minimize overfitting in order to increase the generalizability of model results. One previously studied pruning method is on the basis of sensitivity of the error to each individual weight. In this method, contribution to network behavior is measured explicitly as the sensitivity of the error to each individual weight. Weights below a certain sensitivity cutoff are removed (Mozer 1989). This method is commonly used to prune neural networks (Segee 1991).

The best way to determine if a machine-learning procedure has indeed learned input-output relationships over its domain is to evaluate how well it generalizes to data that it was not trained on (Sietsma and Dow 1991). We trained the model on only the single inputs by removing all of the combination inputs from the truth table (two inputs or more). Non-structure connections of each of the ten original networks were pruned at various cutoffs according to the sensitivity pruning criterion, and the pruned network was then re-trained. During re-training, the structure connections were trained at the same fast learning rate (1) while the non-structure connections were trained at a learning rate that was an order of magnitude lower than that of the fast connections (0.10). The purpose of this was to favor strengthening connections that were known from the experimental literature. Non-structure connections were still trained because they were necessary to produce reasonable agreement with the training set.

We obtained ten fits trained only on single-input data. These ten fits were then pruned at 20 different cutoffs on the basis of sensitivity and re-trained as described in the previous paragraph. The pruned and re-trained weight matrices were then given the drug combinations that were removed from the training set and their outputs were obtained and compared with experimental findings to produce an average RMS error. We found that the original, unpruned weight matrices had an average RMS error of 0.0209 over the untrained drug combination data.

Interestingly, we found that pruning could decrease this average RMS error. We plotted the untrained drug-combination RMS errors versus the number of pruned connections to determine which pruning cutoff on the basis of sensitivity can decrease this RMS value the most. Pruning on the basis of sensitivity at a cutoff of $1x10^{-3}$ pruned 3981 connections on average and produced an average RMS error over the untrained data of 0.0208. All further training sets were trained using the full truth table and pruned at a sensitivity cutoff of $1x10^{-3}$ and retrained using the retraining method described above. This optimized our model's to generalize on acute drug combinations that it was not trained on.

#### S5: Details on Temporal-logic Model-checking Procedure

##### S5.1 Linear Temporal Logic

Linear temporal logic (LTL) is a type of modal temporal logic which allows for reasoning about a sequence or sequences of discrete states evolving in time. It extends propositional and predicate logic by defining additional operators by which one can formally describe relations between temporally linked states. In formal verification, a subfield of computer science, linear temporal logic is commonly used to verify properties of a system, such as liveness (a certain desired property keeps happening) and safety (a certain undesired property never happens). We provide a partial list of LTL operators in the table below.

| **Symbol** | **Operator** | **Description** |
|---|---|---|
| ◊x | Eventually | For at least one state on the subsequent path, x must hold |
| □x | Henceforth | For all states on the subsequent path, x must hold |
| x***U***y | Until | On the subsequent path, x must hold at least until the first state where y is true |
| x \|−> y | Leads to | On the subsequent path, whenever x holds, eventually y must hold |

###### S5.1.1 Notation

For the rest of this section, we will denote configurations of receptor strength adjustments and the steady-state activations of each unit in the network as states. We assume that there are *nadj* adjustable TSCs, *u* network units, and that we explore sequences of at most *d* adjustments. A state can then be written as a *nadj*-dimensional vector **s**, and the corresponding unit activations can be written as a *u*-dimensional vector **u**, all of whose values are in the range (0, 1). The initial state **s0** refers to the state in which no TSC adjustments have occurred, and describes the acute response to a set of drug inputs.

In the most general rendition of a model checking framework, each state may transition to some subset of other states based on a set of rules specific to the model. We choose to model state changes in the biological system by adjusting exactly one of the adjustable TSC strengths by the increment, which is a hyperparameter (here set to 0.50).

A particular sequence of states, for which each state can be reached from its predecessor by a transition, is denoted as a path. The set of all paths originating from some state **s** with depth at most *d* is denoted as **T**(**s**, *d*).

**S5.1.2 Properties**

The structure of this problem yields a number of useful properties. Firstly, the possible TSC strength configurations forms a *nadj*-dimensional ball with *L1*-norm *d*. That is, it contains all of the possible configurations with up to *d* total adjustments from the initial state. It is known that the adjustable TSCs are either inhibitory or excitatory, and it is not biologically plausible for a connection to switch between these two polarities. Our methodology for dealing with issues of polarity and other disqualifying conditions is discussed in more detail in a later section.

In this paper, we are concerned with trajectories of state transitions beginning at **s0**, and therefore evaluate various propositions on interest on **T**(**s0**, *d*). Since the values of linear temporal logic model checks at some state **s** depend only upon the set of possible trajectories beginning at **s**, they may be computed without any knowledge about the trajectory of prior states leading to **s**.

From a biological perspective, each path originating from the initial state corresponds to a possible sequence of TSC strength adaptations in response to drug inputs. The transition rule chosen for this model allows states to transition by incrementing TSC strengths separately. Our model simulates the biological system's ability to concurrently adapt multiple TSC strengths by instead making multiple sequential transitions. However, the model imposes the critical constraint that the TSC strengths in successive states are close to one another. This reflects the property that in the physical system, sensitization and desensitization occur gradually.

In the face of uncertainty about how neuroadaptation occurs in the system, we make the weakest possible assumption, which is that all transitions between two adjacent states which both obey the connection polarity constraint are possible. It is possible that certain TSC strength adjustments do not occur because neuroadaptation only moves the system in a strictly neuroadaptive direction, but neuroadaptation in the monoamine-stress system has not been characterized in sufficient detail to evaluate this possibility. The set of connections between states that we consider is then a superset of the connections that may exist in real life. By considering a system that subsumes the real one, linear temporal logic checks run on our simulation will make conservative predictions. If a property does not hold in the real system, which means that there is at least one trajectory which acts as a counterexample against this property, then a check for this property in our simulation will also find this counterexample and return False. Additionally, if a check for some property in our simulation returns True, then it must hold for all possible trajectories in our simulation, and consequently must also hold for all trajectories in the real system. As a caveat, it is possible for some property to hold for all trajectories of adaptation in the real system, but then return False in our simulation, if there exists some counterexample in the considered superset which is not found in the real system.

##### S5.2 Implementation

###### S5.2.1 Technologies

Linear temporal logic model checks are run using both Maude and Python. Maude is a declarative language implementing rewriting logic, developed primarily by SRI International for formal methods in scientific computing. The primary use case of this language has been as a way to run logical queries on arbitrary systems, which can be programmed and simulated. In contrast with imperative languages like MATLAB, scientists working in declarative languages such as Maude specify the behavior of a program not as a sequence of instructions, but rather as a set of possible states and a set of logical transformations that may be performed on states. This paradigm is well-suited for formal verification tasks in general, since the fundamental task is the simulation of a dynamically changing system, and requires analysis of all possible behaviors, rather than any one predetermined behavior.

Because Maude was written as a logical language for verification, computations in this language compile down to Boolean primitives. Although this yields desirable properties like provable correctness, it is also prohibitively slow for experiments that are computationally intensive. We therefore chose to implement a functionally equivalent system in Python that is also optimized for our specific problem and that takes advantage of its inherent structure. The computation of steady-state unit activations is done in Numpy, which is a matrix arithmetics library for Python, and the rest of the LTL model checking logic is converted into matrix operations in order to use the highly optimized functions provided by Numpy. This implementation is designed to combine the user-friendly and simple syntax of Python with the efficiency of Numpy, which can be orders of magnitude faster than an implementation in pure Python.

###### S5.2.2 Framework Architecture

The framework implemented for model checking in this paper is built to be robust, extensible, and reusable for related problems. Based on the Markov property observed earlier, we may save a substantial amount of computation by making use of the dynamic programming paradigm. This paradigm, which is an offshoot of recursion, avoids recomputing intermediate results in executing a recursive computation by storing each intermediate result, then reusing values stored in this cache whenever needed.

A small proportion of model checks only involve the TSC strength configuration and the unit activations at each state. These are exactly the set of propositions which do not make use of linear temporal logic operators. For this type of query, we simply evaluate the proposition at each possible TSC strength configuration. For instance, the query *(FHT > 0.7)->(Cort < 0.7)* may be evaluated in this manner.

This idea may be extended to the general case of queries involving linear temporal logic operators. Specifically, we may store the truth value of a proposition evaluated at the *d*-th depth, then use those results to compute the truth value of that proposition evaluated at the *d-1*-st depth, and repeat until we obtain the truth value of the proposition evaluated at the *0*-th depth. As a simple example, consider the propositions *Q: Cort < 0.7* and P: Henceforth Q evaluated on **T**(**s0**, 6). This poses the question of whether or not cortisol is below a certain threshold for all trajectories of depth 6 beginning at the initial state. First, this formula must be expressed in a recursive form:

$$P(\mathcal{T}(s,d)) \equiv \begin{cases} Q(s) & d = 0 \\ Q(s) \wedge \bigwedge_{s' \in \mathcal{S}(s)} P(\mathcal{T}(s', d-1)) & d > 0 \end{cases}$$

This new formula can be turned into an iterative algorithm by first enumerating all of the states reachable from **s0** along a trajectory of length 6 and computing whether *Q* holds at those states. Next, compute the value of *P* at each state **s** reachable from **s0** along a trajectory of length 5 by computing whether *Q* holds at that state and whether *P* holds at every depth-6 state reachable from **s**. This proceeds iteratively backwards until the values of *P* at all states for depth 0 are computed. Then, the algorithm returns the value of *P* for **s0** at depth 0.

This construction, unlike the default execution in Maude, takes advantage of the fact that in this system, the truth value of a proposition evaluated at some state does not depend on the trajectory leading to that state. As previously described, we need only refer to results in the most recently-computed depth to determine the next set of results. Therefore, we may delete older intermediate results freely. Because of the size of the configuration space for this experiment, and because we discard all but the most recent intermediate results, it is feasible to keep these values in computer memory. We choose to keep these values in memory rather than on disk for performance, but one could keep results on disk for much larger experiments, or parallelize model checking across multiple processors.

One limitation on the number of model checks, adjustable receptors, and maximum depth in previous experiments was the prohibitive cost of evaluating each path separately in Maude. For our choice of *nadj*=10, and *d*=6, and a simple temporal-logic check, the Maude implementation would perform 64 million comparisons, while our implementation performs approximately 16 million comparisons. The relative performance gains are even more significant for larger values of both depth and adjustable receptors. For a depth of 7 and the same number of adjustable receptors, enumerating each trajectory in the tree separately would require 1.28 billion comparisons, whereas our approach requires approximately 60 million comparisons.

**S5.2.3 Bounds**

As previously noted, certain adaptations can cause adjustable TSCs to have a value that is either of the wrong polarity or exceeds a pre-defined magnitude threshold. In this paper, we reject these adapted states, and remove paths which include these states from the set of trajectories used for model checks.

In all experiments performed on these networks, we fix the maximum absolute TSC strength at 10 for both excitatory and inhibitory connections and the increment at 0.50. The low granularity of TSC strength adjustments was necessary to keep the number of states tractable, but in our experiments, the recurrent neural network displays uniform continuity with respect to model weights. Simulating the effects of finer-grained receptor strength adaptations would not provide much more information than interpolating between our existing results, but would cost significantly more computation time. The maximum TSC strength and increment size were empirically chosen to encourage a variance in the resulting steady-state unit activations that aligns with clinical findings for the diverse chronic effects of certain inputs.

In order to ensure that the linear temporal logic framework is as generalizable as possible, we compute steady-state activations for all states, regardless of whether they obey the specified bounds. Our approach defines a filtering function φ(**s**, **s'**), which encompasses all constraints on the validity of states or state transitions. That is, φ(**s**, **s'**) returns True whenever the transition from **s** to **s'** is a valid one. Here φ merely checks that there is a connection between **s** and **s'** and that adapted TSC strengths for both **s** and **s'** fall within the acceptable range, and False otherwise. However, the idea of a filtering function on transitions is much more generalizable, and gives the user more power in specifying which trajectories are valid. For instance, one could specify that model checks should consider only adjustments that keep a brain region's firing rate above a certain threshold or only adjustments that reduce the difference between the current state's unit activations and baseline unit activations.

Because of this filtering function, our linear temporal logic model checks consider approximately 130,000 states reachable at depth 6, whereas analysis elsewhere in this paper consider approximately 120,000 states, which is the set of remaining states after rejecting states that violate our predefined bounds.

**Supplemental Figure 1: Complete model structure diagram.** This schematic illustrates all of the structure connections of the model. Green, red and blue lines represent excitatory, inhibitory, and trained (determined by the training algorithm) polarities, respectively, between units. Each unit type (drug, receptor, neural region, precursor, metabolite, transmitter, hormone, and enzyme) is represented by a different shape. All units of the same type are arranged in the same row. The file CompleteMSModelStructure.jpeg contains this schematic and can be interactively viewed.

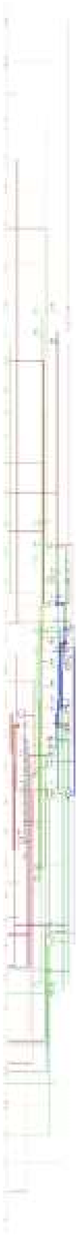

**Supplemental Figure 2: Agreement between desired (i.e., target) and actual outputs is high either after training but before pruning, or after pruning and re-training.** The colormap scales for the desired and actual output plots (A, B, D, and E) are the same, ranging from 0.00 to 0.70. Each row represents the same input pattern and each column represents an output value. The first row represents the no-drug baseline values. The colormap for the absolute differences or errors between the desired and actual outputs (C and F) are between the minimum and maximum absolute difference values for each plot, which is 0.00 to $3.77 \times 10^{-4}$ for (C) and 0.00 to $1.10 \times 10^{-3}$ for (F). Pruning was done to minimize non-structure connections. The RMS error over all training patterns was $2.39 \times 10^{-5}$ for the unpruned network and $5.10 \times 10^{-5}$ for the pruned and re-trained network.

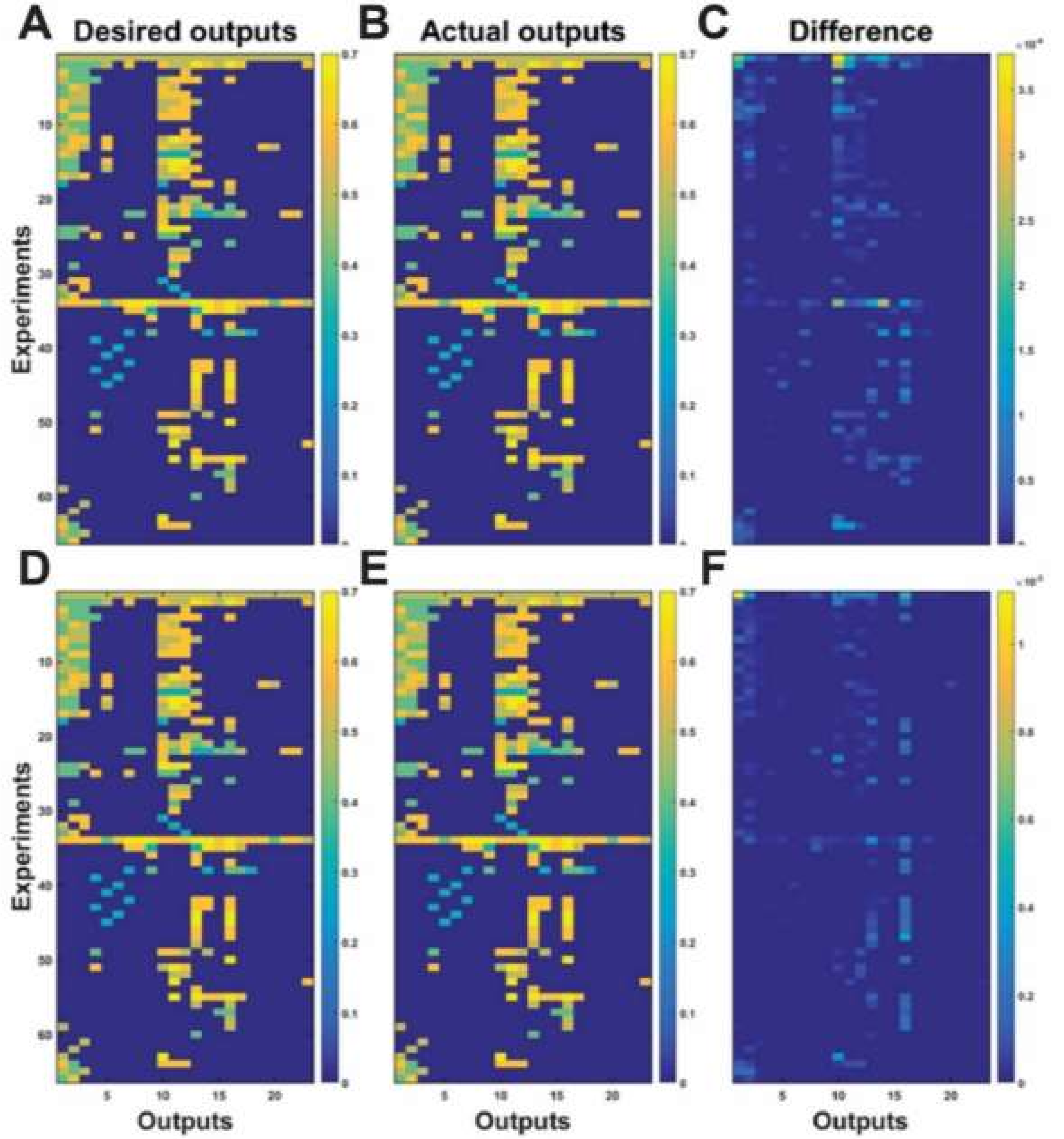

**Supplemental Table 1: Complete model truth-table.** The complete model truth-table can be found Each row of the corresponding table in the Excel file titled CompleteTruthTable.xls represents observed input-output relationships derived from whole organism experiments on the monoaminergic or stress hormone systems. The inputs include drugs, hormones, and lesions applied alone or in combination. Inputs are represented in binary fusion. Most rows incorporate findings compiled from more than one actual experiment. The findings are arranged in input/desired-output pairs and used to train the model. Corresponding references are provided in the Supplemental Text. Abbreviations: amygdala, AM; prefrontal cortex, PFC; hippocampus, hipp; paraventricular nucleus of the hypothalamus, PVN; pituitary gland, PG; adrenal gland, AG; cortisol, CORT; cortocotropin-releasing factor, CRF; arginine vasopressin, AVP; adrenocorticotropin hormone, ACTH; oxytocin, Oxt; melatonin, MT; glutamate, glu; gamma-Aminobutyric acid, GABA; tryptophan, Trp; 5-hydroxytryptophan, 5HTP; galanin, gal; selective serotonin reuptake inhibitor, SSRI.

**Supplemental Table 2: Canonical model weights.** The model has 23 weights that were treated as "canonical weights." The canonical weights are the weights that mediate the known interactions of the monoaminergic nervous system and the stress-steroid system. The training procedure set the lower bound of the canonical weights to absolute value 1. Subsequent analysis revealed that the canonical weights developed the strongest weights and that each network was most sensitive to the canonical weights. They also revealed that the pruning procedure increased the sensitivity of the networks to the canonical weights.

| Number | Weight | Polarity | References |
|---|---|---|---|
| 1 | DR to 5HT | + | (Hensler, Ferry et al. 1994, Monti 2010) |
| 2 | 5HT to 5HT1AR | + | (Davidson and Stamford 1995, Blier, Pineyro et al. 1998) |
| 3 | 5HT1AR to DR | − | (Davidson and Stamford 1995, Azmitia, Gannon et al. 1996) |
| 4 | 5HTT to 5HT | − | (Lesch, Aulakh et al. 1993, Hensler, Ferry et al. 1994) |
| 5 | LC to NE | + | (Grenhoff, Nisell et al. 1993, Samuels and Szabadi 2008) |
| 6 | NE to AR2 | + | (Cedarbaum and Aghajanian 1977, Washburn and Moises 1989) |

| | | | |
|---|---|---|---|
| 7 | AR2 to LC | − | (Cedarbaum and Aghajanian 1977, Washburn and Moises 1989) |
| 8 | NET to NE | − | (Iversen 2000, Bonisch and Bruss 2006) |
| 9 | VTA to DA | + | (Ornstein, Milon et al. 1987, Guiard, El Mansari et al. 2008) |
| 10 | DA to D2R | + | (Benoit-Marand, Borrelli et al. 2001, Perra, Clements et al. 2011) |
| 11 | D2R to VTA | − | (Hall, Sedvall et al. 1994, Perra, Clements et al. 2011, Koyama, Mori et al. 2014) |
| 12 | DAT to DA | − | (Ciliax, Heilman et al. 1995, Iversen 2000) |
| 13 | PVN to CRF | + | (Makara, Stark et al. 1981, Bruhn, Plotsky et al. 1984) |
| 14 | CRF to CRF1R | + | (Van Pett, Viau et al. 2000, Hauger, Risbrough et al. 2006, Holsboer and Ising 2008) |
| 15 | CRF1R to Pituitary Gland | + | (Van Pett, Viau et al. 2000, Nikodemova, Diehl et al. 2002) |
| 16 | Pituitary Gland to ACTH | + | (Makara, Stark et al. 1981, Bruhn, Plotsky et al. 1984) |
| 17 | ACTH to ACTHR | + | (Xia and Wikberg 1996, Papadimitriou and Priftis 2009) |
| 18 | ACTHR to Adrenal Gland | + | (Yang, Koistinaho et al. 1990, Xia and Wikberg 1996, Papadimitriou and Priftis 2009) |
| 19 | Adrenal Gland to CORT | + | (Grant, Forrest et al. 1957, Papadimitriou and Priftis 2009) |

| | | | |
|---|---|---|---|
| 20 | CORT to GCR | + | (Pariante and Miller 2001, Papadimitriou and Priftis 2009) |
| 21 | GCR to Adrenal Gland | − | (Loose, Do et al. 1980, Kalinyak, Dorin et al. 1987) |
| 22 | GCR to Pituitary Gland | − | (Morimoto, Morita et al. 1996, Ozawa, Ito et al. 1999) |
| 23 | GCR to PVN | − | (Morimoto, Morita et al. 1996, Ozawa, Ito et al. 1999) |

**Supplemental Table 3: Adjustable TSCs.** Neuroadaptation was simulated by allowing the networks to adjust the strengths of the weights that are known to adjust under the conditions of the experiments from which the truth table was derived. The adjustable TSCs were a subset of the canonical weights. Configurations of adjusted TSCs that resulted in a reduction in network imbalance by bringing the most heavily trained brain regions (DR, LC, VTA, PVN) back toward their baseline values were considered adapted.

| Number | Adjustable TSC | Polarity | References |
|---|---|---|---|
| 1 | 5HT1AR to DR | − | (Blier and de Montigny 1987, Szabo and Blier 2001, El Mansari, Ghanbari et al. 2008, Ghanbari, El Mansari et al. 2010, Rozeske, Evans et al. 2011) |
| 2 | AR2 to LC | − | (Szabo and Blier 2002, El Mansari, Ghanbari et al. 2008) |
| 3 | D2R to VTA | − | (Chernoloz, El Mansari et al. 2009, Katz, Guiard et al. 2010, |

| | | | |
|---|---|---|---|
| | | | Madhavan, Argilli et al. 2013) |
| 4 | 5HTT to 5HT | − | (Lesch, Aulakh et al. 1993, Benmansour, Cecchi et al. 1999, Lau, Horschitz et al. 2008) |
| 5 | NET to NE | − | (Hebert, Habimana et al. 2001, Miner, Jedema et al. 2006, Pietrzak, Gallezot et al. 2013) |
| 6 | DAT to DA | − | (Neumeister, Willeit et al. 2001, Brunswick, Amsterdam et al. 2003, Kugaya, Seneca et al. 2003, Yang, Yeh et al. 2008) |
| 7 | GCR to PVN | − | (Pariante and Miller 2001, Barden 2004, Ladd, Huot et al. 2004) |
| 8 | GCR to Pituitary Gland | − | (Barden 2004, Ladd, Huot et al. 2004) |
| 9 | GCR to Adrenal Gland | − | (Barden 2004, Ladd, Huot et al. 2004) |
| 10 | CRF1R to Pituitary Gland | + | (Nikodemova, Diehl et al. 2002, Kageyama, Hanada et al. 2006) |
